# Introducing ARTiMiS: A low-cost flow imaging microscope for phytoplankton monitoring in engineered and natural environments

**DOI:** 10.1101/2024.02.27.582145

**Authors:** Benjamin Gincley, Farhan Khan, Elaine Hartnett, Autumn Fisher, Ameet J. Pinto

## Abstract

Manual microscopy is the gold standard for phytoplankton monitoring in diverse engineered and natural environments. However, it is both labor-intensive and requires specialized training for accuracy and consistency, and therefore difficult to implement on a routine basis without significant time investment. Automation can reduce this burden by simplifying the measurement to a single indicator (e.g., chlorophyll fluorescence) measurable by a probe, or by processing samples on an automated cytometer for more granular information. The cost of commercially available flow imaging cytometers, however, poses a steep financial barrier to adoption. To overcome these labor and cost barriers, we developed ARTiMiS: the Autonomous Real-Time Microbial ‘Scope. The ARTiMiS is a low-cost flow imaging microscopy-based platform with onboard software capable of providing species-level quantitation of phytoplankton communities in real-time. ARTiMiS leverages novel multi-modal imaging and onboard machine learning-based data processing that is currently optimized for a curated and expandable database of industrially relevant microalgae. We demonstrate its operational limits, performance in identification of laboratory-cultivated microalgae, and potential for continuous monitoring of complex microalgal communities in full-scale cultivation systems.

**Synopsis:** We introduce a platform for low-cost real-time imaging monitoring of phytoplankton and demonstrate its utility in real-time monitoring of laboratory- and full-scale microalgal cultivation systems.

## Introduction

Biotechnologies that utilize phytoplankton (i.e., microalgae), including both pure cultures and mixed communities, are becoming increasingly important for nutrient removal^1^, resource recovery^2^, carbon capture^3^, and for the production of petroleum-replacing feedstocks for industrial use-cases^4^. Rapid monitoring is critical to improve process stability as real-time sensing can enable the use of closed-loop engineering controls. While real-time sensors for water chemistry parameters (e.g., pH, temperature, etc.) are widely used for process control and monitoring, detailed measurement of the biological characteristics of such systems is less commonplace. Devices that measure optical density or fluorescence provide results in real time, but report proxy measurements for total biomass. While offline analytical techniques have been developed to improve accuracy^5^, these devices ultimately lack the single-cell resolution necessary to provide detailed information on taxonomic identity and morphological population statistics, such as changes in cell size over time, that may be critical for optimal process monitoring and control.

Manual microscopy has been the primary method for phytoplankton identification and enumeration^6^ until the development of automated microscopy instruments^7^, and still remains the gold standard approach^8^. For accurate taxonomic identification in both manual and automated approaches, a trained taxonomist is required. Automated microscopes leverage advances in digital camera technology and computer vision to reduce the time-consuming and labor-intensive work of particle identification by creating a digital collection of images that can be annotated asynchronously. The annotation process requires a curated database, and online database communities such as EcoTaxa^9^ crowdsource the task of accurately identifying microscopic particles imaged from environmental systems. One major criticism of automated digital microscopy is the lack of sample interactivity: digital images cannot be re-focused or examined at multiple magnifications, unlike live samples on a benchtop microscope. Nonetheless, there has been a consistent decline in the number of professional taxonomists over recent decades^10^. As a result, if phytoplankton monitoring programs lack the time or expertise to routinely perform identification themselves, they must rely on shipping samples to a dedicated phycology lab. The net result is a barrier to implementing proactive measures, informed by microalgal monitoring data, that may mitigate or prevent process upsets or losses in performance^11^.

To meet the need to identify microalgae in a more timely manner, a number of flow imaging microscopes/cytometers have been developed since the late 1990s, among which FlowCam^12^ and the Imaging Flow CytoBot^13^ remain two of the more commonly cited commercial solutions. However, these instruments are often price-prohibitive for prospective end users. This has driven the scientific community to develop alternative low-cost solutions, such as LudusScope^14^, SAMSON^15^, HABscope^16^, PlanktoScope^17^, and other contemporary developments. Each aspires, at least in part, to reduce the capital barriers to entry. This “next generation” of low-cost digital microscopy systems is built upon recent technical and economic advancements in microfluidics, computing, and digital camera technology, the latter of which are in part a byproduct of the smartphone industry; in fact, several projects are designed to use a smartphone as the imaging, computing, and data transfer component (e.g., HABscope). One major drawback among advances in next-generation flow imaging microscopy is the apparent lack of a complete, “batteries-included” solution for professional users who lack the engineering or phycology resources required to make full use of such projects in the literature. Many of the aforementioned devices are targeted at citizen science programs and are freely available as open software/hardware products. However, end-users are often responsible for sourcing components for, and assembly of, the microscope systems. While some projects require users to perform taxonomic analysis themselves, others such as PlanktoScope provide functionality to upload data to the EcoTaxa database for community annotation. In all cases, the process of effectively managing the requisite data workflows to transform raw image data into refined, actionable results imposes a technical barrier to adoption; the cost of switching from existing solutions (i.e., shipping samples to a commercial phycology laboratory) can make adoption of such projects less attractive for teams looking for a turnkey solution.

To address this, we have developed ARTiMiS, the Autonomous Real-Time Microbial (micro)’Scope, a low-cost flow imaging microscopy platform with companion digital image processing and phytoplankton identification software for real-time, on-premises results. The ARTiMiS is designed to mitigate both the financial and technical barriers to adoption normally associated with digital flow imaging microscopy technologies by providing an end-to-end automated platform that is capable of accurately estimating phytoplankton identity and abundance in engineered and natural systems.

## Materials and Methods

### Design and Construction of the Fluidic, Optical, Actuation, and Automation Systems

The instrument was designed to incorporate off-the-shelf components where possible, with custom solutions implemented to bind components and the full system together as required. The device housing was 3D printed (**Figure 1A**), and a printed circuit board (PCB) was designed to route connections between the power supply, motor controllers, actuators, and the single-board computer. The system was designed to be logically controlled by Raspberry Pi 4 Model B and required a 12V 3A DC power supply. User control was facilitated via a Graphical User Interface (**Figure 1B**). The optical system consisted of a programmable LED array (UnicornHAT HD, Pimoroni® PIM273) serving as the light source (**Figure 1C**), which allowed both bright and dark field illumination of each specimen (**Figure 1D**). Magnification was provided by two M12-threaded board lenses in a reverse lens configuration, as reported in contemporary designs (Octopi^18^, PlanktoScope^17^). An 8MP Raspberry Pi Camera V2 (Sony IMX219 sensor, stock lens removed) served as the camera sensor. The fluidic subsystem centered around a commercially available microfluidic chip with a 200 µm deep flow channel, which was flanked by two isolation valves for stop-flow operation to allow particles to settle within the field of view. At benchtop scale, samples were recirculated using a low-cost peristaltic pump marketed for aquarium applications. Focusing was controlled via a stepper motor linear actuator. A complete list of materials (**Table S1**) and design methodology (**Text S1**) is provided in supplementary materials.

**Figure 1:**
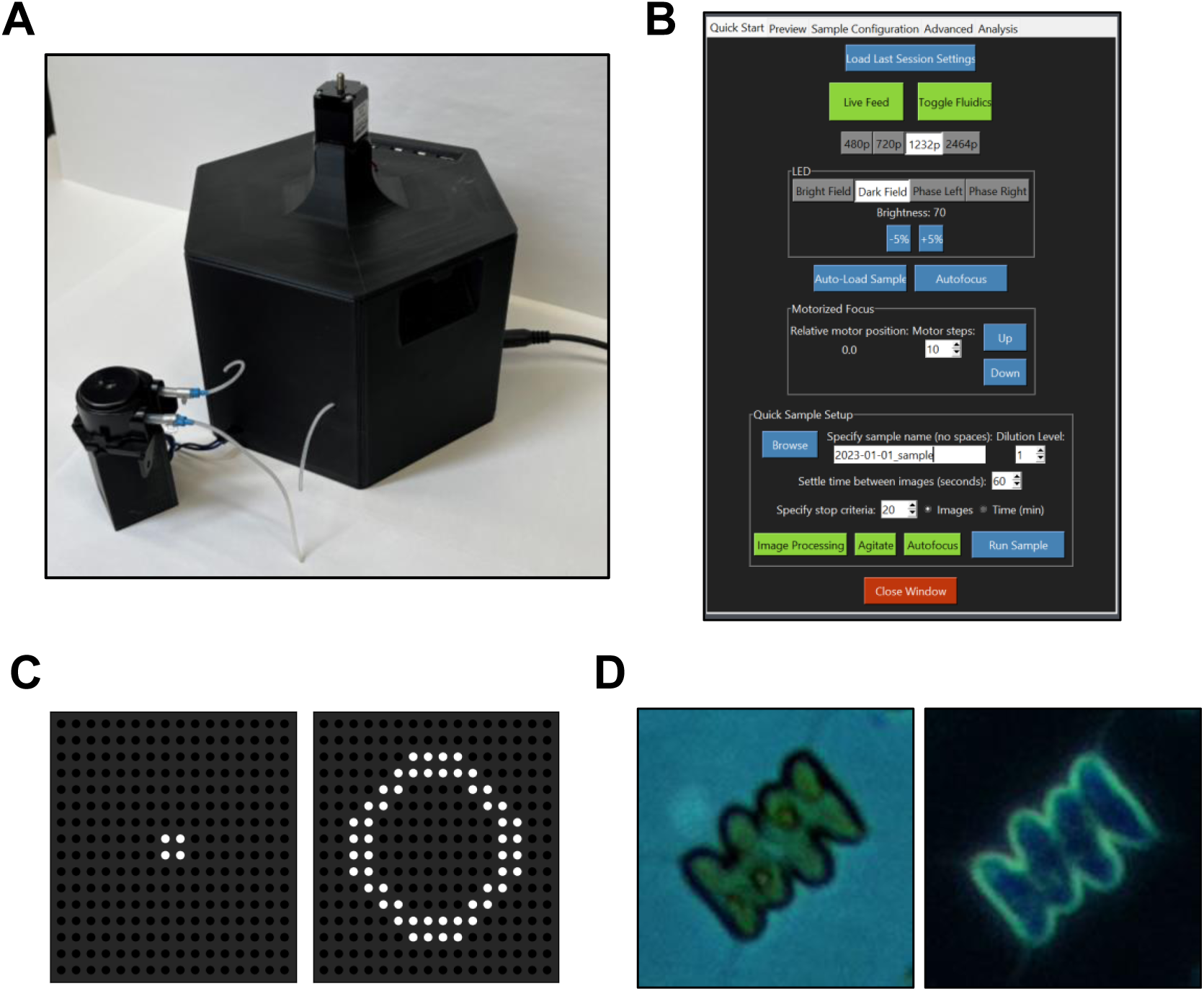
ARTiMiS (A) was designed as a compact, portable device for both benchtop and in-field use. A graphical user interface (B), enabled both configuring sample processing settings and manual instrument control, including a live camera view, and was accessible by both wired and remote connections. A key functional element was the programmable LED array (C), which enabled capture of brightfield and darkfield images of the same field of view (D). This provided complementary images containing intracellular detail and high foreground contrast of microorganisms, such as *Scenedesmus* sp. (shown).

### Assessing Optical Resolution and Magnification

A USAF resolution test target (Edmund Optics Inc.) was centered in the field of view and then the focus was manually adjusted. It was less reliable to focus the image based on the smallest elements, Group 9 (**Figure S1A**), due to pixel binning from the camera sensor to the camera preview display. Instead, adjustment was performed until the maximum apparent contrast between line pairs in Group 8 could be observed. For quantitative image analysis, multiple views of the same elements were captured in multiple locations about the field of view to account for the impact of radial position on color and resolution^19^. Six total views were captured with both the 16 mm focal length (FL) and 25 mm FL lenses, positioning the smallest groups near each of the four corners of the image and two views at the center. Each image was captured using brightfield illumination at the same illumination intensity. Blank images were also captured using the full-frame open glass cutout of the test target (positive blank), as well as a fully obstructed view centered on the test target opaque coating (negative blank), for normalization. From the test target data, the resolution of the imaging system was calculated from the smallest separable line pair element as described by Pollina et al (2022) ^17^.

### Object Detection Approach and Performance Assessment

Image pre-processing was required to normalize luminance and reduce noise prior to obtaining regions of interest (ROIs) during object detection (**Text S2**). Bounding box coordinates for each ROI were identified using foreground segmentation, then expanded by approximately 8µm in post-processing to provide additional visual context to aid the observer in downstream identification (i.e., Segmenter approach). An alternate particle detection algorithm based on local intensity maxima detection was developed to accurately identify uniformly shaped particles while significantly reducing computational time (i.e., Fast Detector approach). Both object detection algorithms were assessed for accuracy and computation speed. Four different sample types were used to provide a holistic representation of expected performance under varying conditions: pure culture and mixed microalgal community samples at medium (approximately 5×10^5^ cells/mL) and high (approximately 5×10^6^ cells/mL) cell densities. Replicate (n=6) runs were performed for statistical analysis. Object detection approaches were scored manually, with ‘counting chamber’-like gridlines digitally drawn on each image to simulate the hemocytometer counting method. Each widefield image was divided into 12 counting squares approximately 800×800 pixels (160 µm) in size. For each counting cell, all valid objects were counted as ‘total’ objects. Object detector-supplied ROIs were counted as ‘perfect crops’ having no defects (i.e., true positive or correct). Each ROI that encapsulated <90% of the targeted valid object was counted as a ‘split object’ ROI. Each ROI, encapsulating a primary valid object and at least 50% of one or more additional objects, was counted as a ‘multi-object’ ROI (i.e., merges). All ROIs not containing any valid object were counted as false positives. All objects not included in any ROI were counted as false negatives. Computational speed was assessed on a Raspberry Pi single board computer to represent in situ performance, as well as on a desktop computer processor to represent asynchronous sample analysis. The Raspberry Pi 4 Model B central processing unit was a Broadcom BCM2711 (4 cores, 1.8GHz) with 4 GB RAM. The desktop computer processor used was an AMD Ryzen 9 5900X (12 cores, 3.7GHz) with 32 GB RAM, representing a “best case” hardware scenario. To measure computation speed, a representative sample image was pre-loaded into memory, then full pipeline processing time was measured.

### Determining Limit of Blank, Detection, and Quantitation

Limit of Blank (LoB), Limit of Detection (LoD), and Limit of Quantitation (LoQ) were measured by adapting protocols described previously^20–22^ and the protocol is described in detail in **Text S3** in the supplementary material. Briefly, LoB was measured using blank samples, and LoD and LoQ were measured using a serial dilution of laboratory-cultivated *Chlorella sorokiniana* (UTEX1602), performed as a parabolic dilution series as described by Hubaux and Vos^22^. Measurements were made in parallel on ARTiMiS and a flow cytometer (CytoFLEX, Beckman Coulter) with paired replicates run simultaneously on each instrument.

### Generation and Curation of Training Data for Particle Classification

Training data for particle classification was generated using a range of abiotic and biotic particle types obtained from laboratory-specific and field-relevant conditions. Specifically, Polystyrene Polybead® microspheres (Polysciences, Inc.) of six diameters ranging from 1 to 15 μm were used as a sample material of known physical dimensions for instrument calibration. Two strains of green microalgae were cultured in a laboratory setting to provide biological sample material with known taxonomic ground truth. *Chlorella sorokiniana* (UTEX1602) and *Scenedesmus obliquus* (UTEX393) were procured from the University of Texas, Austin algae culture collection (UTEX) and grown in Bold’s 1NV medium in batch cultures at 25°C under a 12h:12h day-night cycle at a PAR value of 80 µmol photon/m^2^s. Field-relevant samples were obtained from a full-scale EcoRecover process at the Roberts Water Treatment Plant (Village of Roberts, Wisconsin, USA); a mixed microalgal cultivation system designed and operated for phosphorus removal^23^. Samples were collected daily and immediately stored at 4°C refrigeration, then shipped overnight to Georgia Tech in 15 mL conical tubes for processing.

Image curation was performed by a human annotator as follows: microspheres were imaged in individual samples by size, and resulting ROIs were binned by the degree of focus (DoF, in- or out-of-focus) to create libraries of known size class and human-annotated DoF class. Particle features were extracted using methods from Python libraries Scikit-Image (v0.19.2), OpenCV (v4.6.0), and a bespoke feature extraction library (**Table S2**). Monospecific laboratory-cultivated microalgae ROIs were annotated as in-focus, out-of-focus, ROIs containing multiple distinct individual cells, and ROIs containing non-algal debris particles as determined by a human annotator. Only in-focus libraries were used in taxonomic microalgae classification analysis. Libraries of representative examples of the dominant taxonomic groups identified in EcoRecover samples were curated from representative samples from select time windows between November 2021 and August 2022. Training data libraries consisted of 1,500-2,000 unique objects per class. The “unknown” class was constructed from images of out-of-focus or otherwise indistinguishable objects as evaluated by a human annotator.

### Image Classification Model Construction, Training, and Validation

All machine learning workflows were implemented in Python (v3.9.12). Annotated datasets, as described above, were divided into “train” and “test” subsets using a “train-test split” subroutine (Scikit-Learn, v1.1.1) at a ratio of 85:15. The sample sizes of each test population are indicated in respective confusion matrix plots, e.g., Class A (n=150) describes a total library size of 1000 unique objects, 150 of which comprised the test dataset. Random forest (Scikit-Learn, v1.1.1) algorithm parameters were optimized using a random grid search optimization step to select highest-scoring parameter combinations, optimized for F1 score, prior to final model creation and training. Example hyperparameters for the model used in Random Forest model (see **Figure 5B**) were as follows: {n_estimators: 250, min_samples_split: 5, min_samples_leaf: 2, max_features: sqrt, max_depth: 30, bootstrap: False}. Feature importance was ranked based on mean decrease in impurity, computed as the mean, standard deviation of impurity decrease accumulation in each tree (Scikit-Learn, v1.1.1). A convolutional neural network (CNN) deep learning model (TensorFlow, v2.5.0) of the architecture layout described by Zhou et al (2017) ^24^ was trained from randomized initial weights using annotated microalgae image data. After creating the holdout “test” data subset, the remaining training data was augmented eightfold using all combinations of 90-degree rotations and mirrors of each ROI. Models were trained for between 12-24 epochs in three cross-validation batches, which randomly resampled training and validation data subsets. Model architecture and hyperparameters were as follows: INPUT-32CONV(5×5)-MP(2×2)-16CONV(5×5)-MP(2×2)-8CONV(3×3)-MP(2×2)-16FC-*N*FC where *N* describes the number of output classes, CONV is 2D Convolution layer, MP is Max Pooling layer, FC is fully-connected (dense) layer; {output activation: softmax, optimizer: Adam, learning_rate: 0.01, loss: categorical_crossentropy}. The final model variant was selected as having the highest overall accuracy while minimizing inter-class accuracy variability among 10 random initializations.

## Results and Discussion

### The ARTiMiS Demonstrates Micron-Level Resolution

The resolution of ARTiMiS was determined using a resolution test target (**Figure S1A**). Resolution was estimated based on the smallest resolvable group of line pairs that could be observed through the instrument. Calculating resolution from the smallest line pair (Group 9 Element 3, **Figure S1B**) the empirical resolution was determined to be 1.55 µm. Magnification can be theoretically estimated by the ratio of focal lengths between the objective and tube lenses, though true empirical magnification must be measured with calibration samples (**Figure S1C**). Empirical magnification was determined using the test target manufacturer’s prescribed line width (**Figure S1D**) and the width of each sensor-side pixel (1.12 µm). The 25mm FL objective lens variant, with a theoretical magnification of 8.9X, measured an empirical magnification of 7.8X. The 16mm FL objective lens variant, with a theoretical magnification of 5.7X, measured an empirical magnification of 5.0X. While magnification appeared nominally low for this microscope system, it is important to note that the image is projected onto a camera sensor with small pixels (1.12 µm); thus a 2 µm object magnified at 5X appears as a 10 µm image sensor-side, which spans approximately 10 pixels. Thus, a sufficient Nyquist sampling frequency can be achieved by this system even for single bacterium cells under high contrast (i.e., dark field).

### Object Detection Approaches were Tuned to Sample Matrix Types

We designed an object detection computer vision pipeline to allow the ARTiMiS to automate particle identification and counting. Two different object detection algorithms were developed to meet divergent requirements (**Figure S3**). First, a less computationally intensive algorithm was optimized to detect objects of uniform morphology, such as monospecific unicellular microalgae (e.g., *Chlorella*). A second object detection algorithm was developed to accommodate samples containing particles of varying morphologies, e.g., from mixed microalgal communities or environmental samples. This method required more extensive image pre-processing to yield accurate results and was thus more computationally intensive. These two algorithms are referred to as the “Fast Detector” and the “Segmenter,” respectively, as the latter utilizes segmentation-based image processing techniques, and the former is significantly faster at runtime (**Table S3**).

Object detection algorithms output a “region of interest” (ROI), a cropped region from the larger original image that contains an object to be further examined. An ideal ROI would contain a single, visually intact, and analytically relevant particle. To evaluate object detection algorithm performance, both correct and incorrect detections were quantified. To disambiguate incorrect detections, a more comprehensive set of metrics^25^ was evaluated for each object detection algorithm rather than an all-encompassing metric such as Intersection over Union (IoU)^26^ or mean Average Precision (mAP). Though these condensed scores provide singular, universal values to compare competing methods, they do not fully convey nuanced differences in the types of errors incurred by different algorithms. For instance, IoU relies on pixel-perfect matches between algorithm prediction and manually annotated ground truth to produce a score of 100%. We determined that such scores would not be directly comparable between the two algorithms described here, as one algorithm allows for flexible bounding box geometries, while the other produces fixed-shape bounding boxes; this makes truly fair evaluation for the purpose of comparison impractical.

To circumvent these limitations, we classified each ROI into five classes: true positive (i.e., correct), false positive, false negative, merge, and split; the latter four representing failure types (see **Materials and Methods**). Among failure classes, merge ROIs represented a “less costly” error provided they can be further refined into separate objects in re-analysis. Similarly, with additional post-processing, false positives could be filtered out as noise and discarded. Split errors bore a higher cost to correct, requiring not only a means to be accurately identified but also a method to reconstruct split objects. False negatives were the highest-cost errors to incur, as these objects would be lost entirely. Two different sample types, a monospecific *Chlorella sorokiniana* culture and field samples from a mixed microalgal community (**Figure S3**), were selected to demonstrate the differences in algorithm performance across different sample conditions. Each matrix was assessed at two cell densities (∼5×10^5^ particles/mL and ∼5×10^6^ particles/mL), referred to as “medium density” and “high density,” respectively (**Figure S4**), to quantify the impact of visual crowding on detector performance.

For monoculture samples at medium cell density (**Figure 2A**), the Fast Detector achieved more correct ROIs and fewer erroneous ROIs compared with the Segmenter in significantly less computation time (**Table S3**). At higher cell densities (**Figure 2B**), correct ROIs decreased, though this was largely accounted for in the less-expensive merge errors for Fast Detector. As false negatives and positives remained low at both densities compared to Segmenter, the Fast Detector was considered optimal for pure culture algal samples. Low Fast Detector accuracy (<70%) on the mixed microalgal community sample type (**Figure 2C, 2D**), however, made evident the necessity of the Segmenter algorithm. Importantly, a majority of Fast Detector errors were the more-expensive false negative and split errors. In contrast, the Segmenter achieved a high yield of correct ROIs (>90%) and few to no false negatives or splits, at the expense of slower runtime (**Table S3**). Thus, the Fast Detector algorithm was thereafter used for samples of homogenous cell material, e.g., monoculture single-celled microalgae, and the Segmenter was used for all samples containing multiple species. The Fast Detector is well-suited to quickly enumerate samples such as monospecific cultures in scenarios where results may be needed in true real-time (available within seconds). In contrast, the Segmenter yields accurate ROIs from mixed community samples – more representative of open systems – which would likely be worth the comparative slight delay in results (available within minutes). Nonetheless, both algorithms automate particle counting, removing the burden of manual enumeration. Furthermore, archiving digital images provides a digital record of each sample for later review or analysis, after the biological sample has expired.

**Figure 2:**
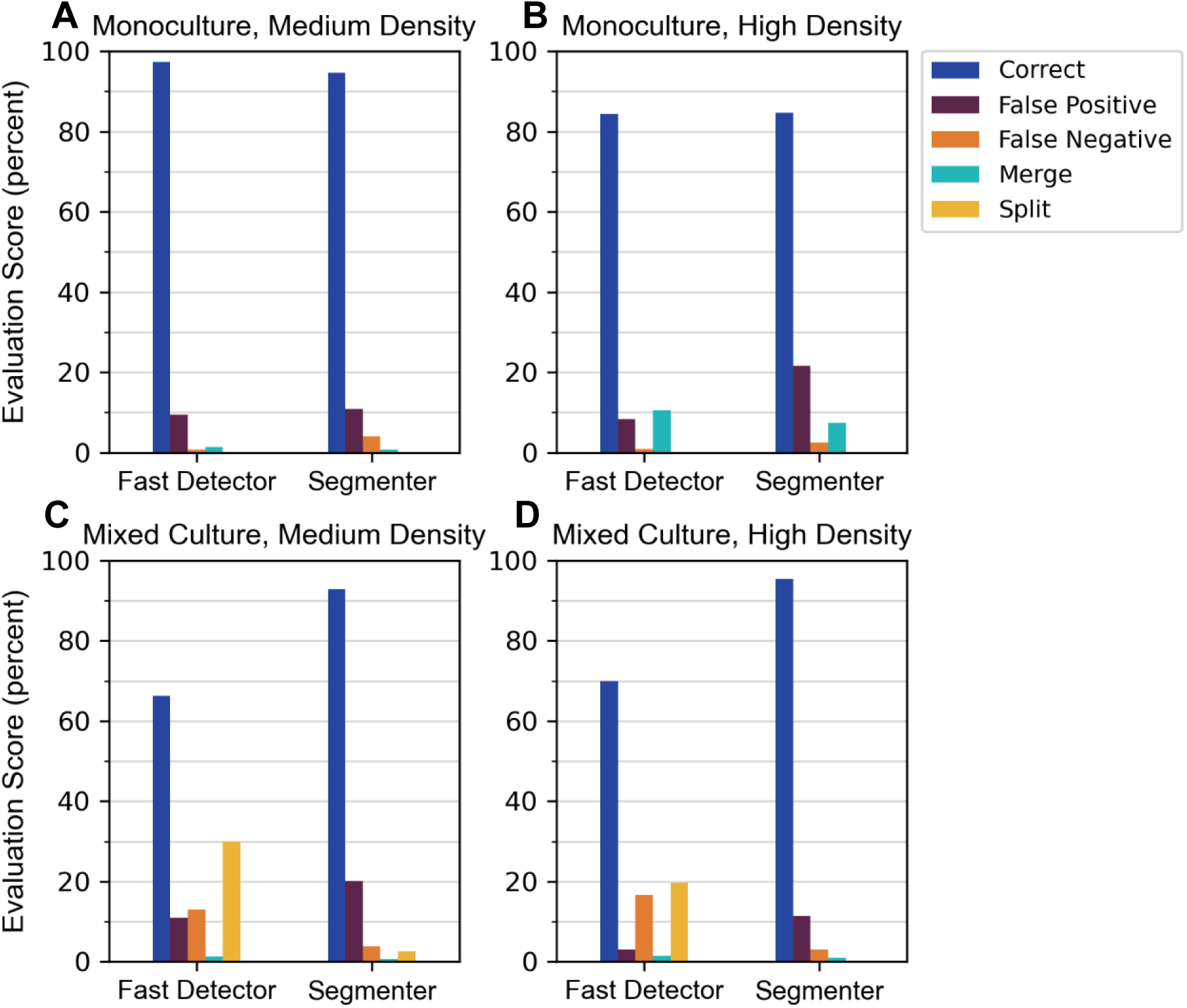
Object detection algorithms “Fast Detector” and “Segmenter” were scored by performance across varying sample types to assess the frequency of multiple types of error under different operational conditions. Samples containing monospecific laboratory cultures of Chlorella sorokiniana (A, B) were compared to samples from a mixed microalgal community (C, D) at both medium (A, C) and high (B, D) cell densities. Error types include false positives (ROIs with no objects), false negatives (missed objects), merges (single ROIs containing multiple objects), and splits (single objects divided by multiple ROIs). All counts are normalized to the total number of valid objects to calculate a percentage score.

### Experiments with Synthetic Samples Highlighted the Impact of Out-of-Focus Particles on Accurate Quantitation and Classification

Approaches to automate classification range from curated thresholds applied on specific features^27^ to multivariate data-driven techniques that determine a “fingerprint”^28^ for each class without requiring *a priori* feature knowledge. To this end, while machine learning is an effective tool, its accuracy is highly dependent on intra- and inter-class variability. To maintain accuracy for complex samples, there is often a requirement to increase the complexity of the classification approach to match. To assess the baseline classification potential of ARTiMiS, synthetic polystyrene microspheres manufactured to specific diameters were used. Each size class was processed on ARTiMiS using the Fast Detector algorithm. Geometric features of particles were measured (**Table S2**) and were used to train a random forest machine learning model to differentiate between each size class. The ten most influential features for class differentiation were ranked are shown in **Figure 3A**. Among these features, a majority related to object size characteristics; this indicated ARTiMiS was precisely measuring objects. Classes with greater size differences (e.g., 10 and 15 µm) consistently exhibited clear separation (**Figure 3B**). Of note regarding accuracy, features were measured from the dark field view by which microspheres appear as rings of their inner refractive diameter (**Figure S5**); thus, their apparent diameter represents this attribute. Likewise, ARTiMiS’ point spread function (**Figure S6**) represents the narrowest measurable diameter, flattening apparent diameters at the 1 to 2 µm end of the range. After training the model on a subset of the complete data set (the “training” set), the model made predictions on unseen data (the “test” set) for evaluation (**Figure 3C**). Among the 127 test images of 4.5 µm particles, 98.4% were correctly labeled by the model with the remaining fraction misclassified as belonging to the 6 µm class. A similar degree of accuracy and inter-class confusion was observed among the 1 µm and 2 µm bead classes. A cause for this misclassification was observable in **Figure 3B**, where regions corresponding to each class overlap for smaller-sized bead classes. This could be attributed to visual noise and the effect of object focus, where objects may appear larger than their true size (**Figure 3D**, **Figure S5**).

**Figure 3:**
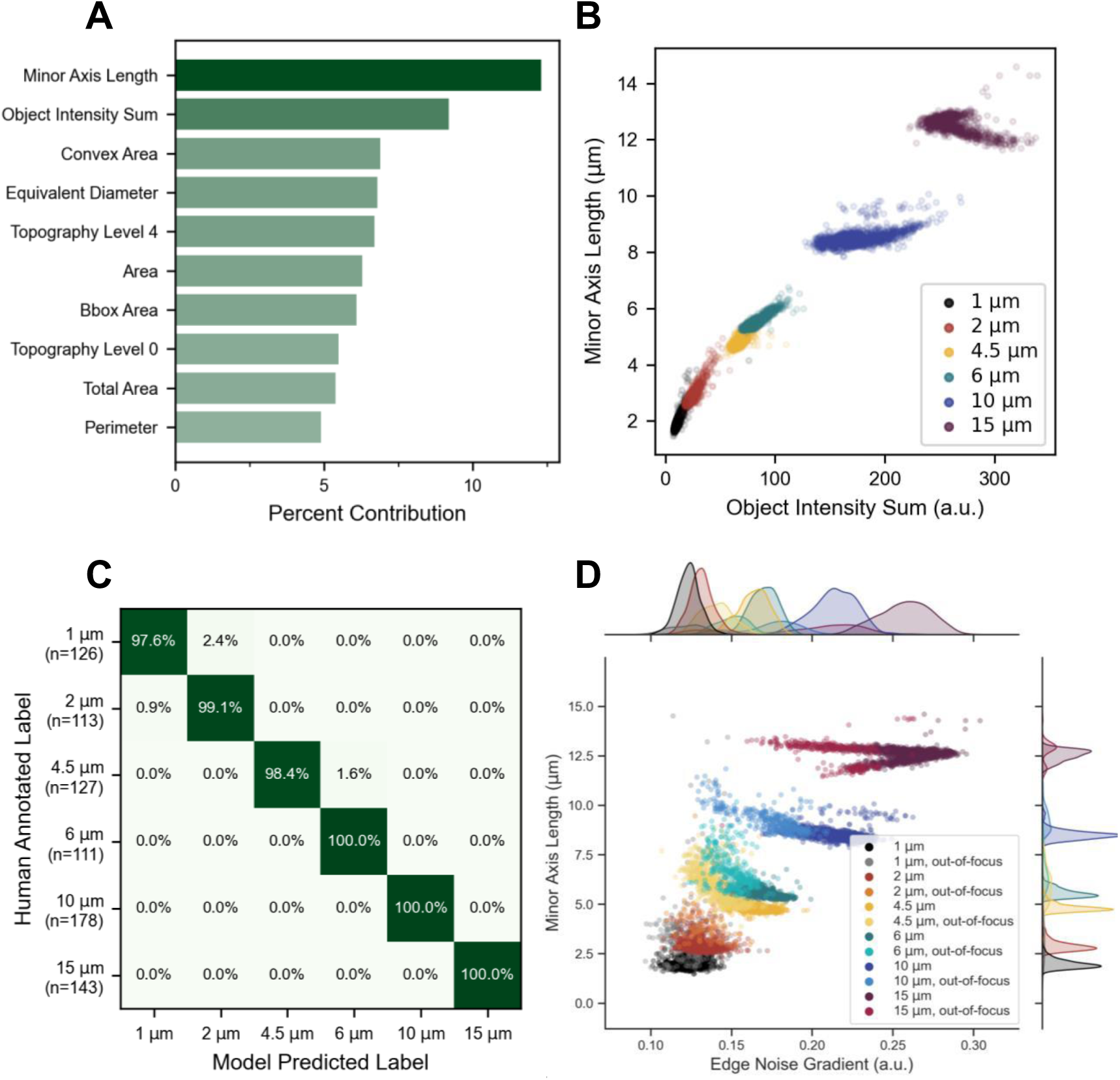
A random forest classifier was trained to distinguish between in-focus samples of each size class, and the importance of individual morphological measurements (features) were ranked (A) by percent contribution for class separation in the decision trees. Dominant features predominantly directly or indirectly described object size characteristics. Examination of the top two features (B) demonstrated threshold separability of classes, especially at larger sizes. Minor Axis Length describes object diameter, while Object Intensity Sum describes total luminance in a darkfield image. With this “rectangular” feature space, a random forest classification model was a well-suited classifier type and could be trained with very high (99.2%) overall accuracy (C) when evaluated on an unseen test dataset: on-diagonal values represent correct model predictions, off-diagonal values represent misclassification rates per class. Number of samples in the test dataset is indicated on annotation axis. A primary source of misclassification in test data and live samples was out-of-focus particles (D), which can be partially distinguished from their in-focus counterpart class with the Edge Noise Gradient feature. Marginal axes depict population distributions from annotated collections of in- and out-of-focus training data, with individual objects shown as scatter points.

### ARTiMiS Detection Limits and Dynamic Range were Comparable to Conventional Flow Cytometry

A parabolic dilution series^22^ of *Chlorella sorokiniana* culture was performed to assess the accuracy of labeled particle counting across a range of concentrations. Each sample point was processed on both ARTiMiS and a flow cytometer (CytoFLEX, Beckman Coulter Life Sciences), a gold standard in cell counting to serve as a ground truth for comparison. “Gating,” i.e., excluding objects outside a specific threshold, is conventionally used to filter non-target events in flow cytometric analysis to improve specificity. Filtering sources of quantification noise, including out-of-focus particles and debris, was likewise a necessary step for ARTiMiS data post-processing (**Figure 4, Figure S7**). As highlighted in the analysis of object detection, ROIs can contain more than one valid object (merge ROIs). These ROIs constituted a larger fraction of the dataset as sample concentration increased. Therefore, four distinct classes: high-quality (in-focus) cells, out-of-focus cells, debris particles, and multi-particle (merge) ROIs; were quantified separately. Object classification was utilized to estimate the abundance of each of these classes. In the same manner as described previously, sample data were manually annotated to produce labeled libraries containing representative images of each class, and this dataset was used to train a classification model (**Figure S7**) that could then be used to predict the abundance of each class in dilution series samples (**Figure 4**).

**Figure 4:**
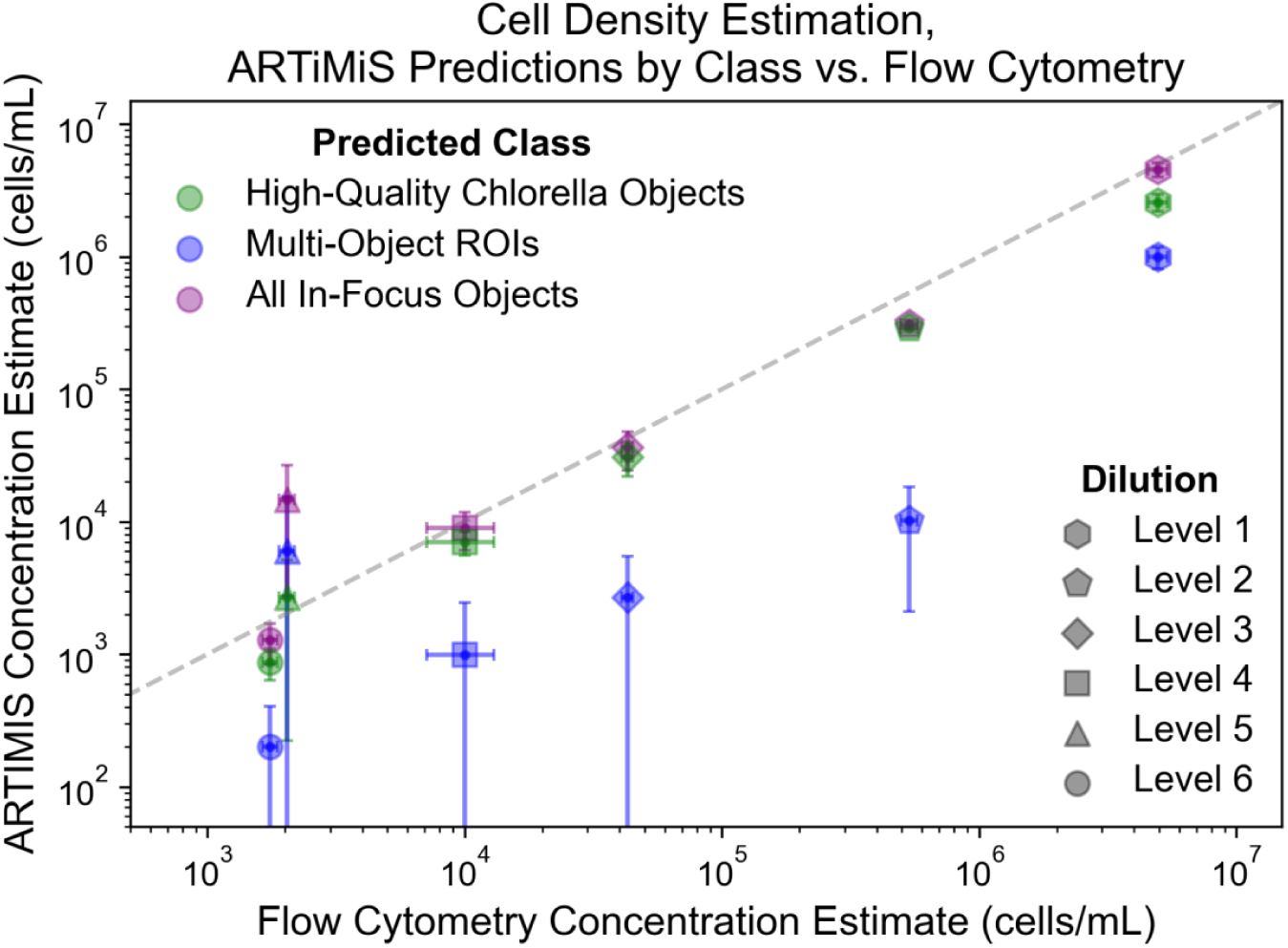
Quantitative comparison of enumeration estimates from both flow cytometry and ARTiMiS compared, describing samples across six levels of dilution. Cumulative dilution levels are as follows: Level 1: 12x, Level 2: 120x, Level 3: 960x, Level 4: 5760x, Level 5: 23040x, Level 6: 46080x (dilution series: 12x, 10x, 8x, 6x, 4x, 2x).

**Figure 5:**
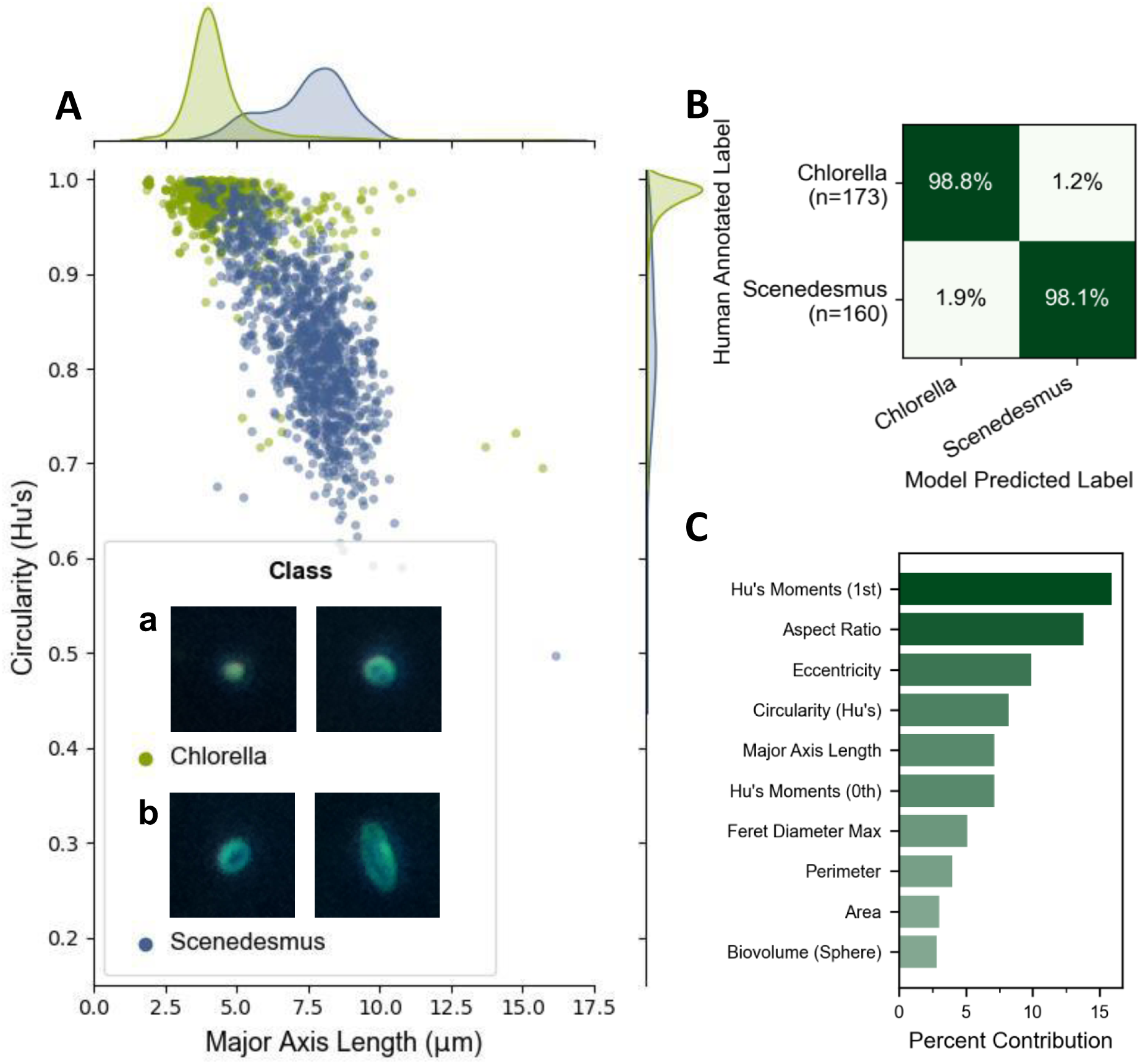
Differentiating between Chlorella (A, inset a) and Scenedesmus (A, inset b) cells. Population distributions demonstrate overlapping feature regions with long tails (A). Representative images of each class, demonstrating intra-class morphological diversity and inter-class similarity for some individuals, are shown in the legend inset. A random forest classifier trained using all available features achieves 98.5% overall accuracy (B). Features informing population separation decision boundaries relate primarily to circularity/ellipsoid-ness (C).

For intermediate concentration levels (1 x 10^4^ to 1 x 10^6^ cells/mL), ARTiMiS estimates were equivalent to concentrations reported by flow cytometry (**Figure 4**). At higher concentrations (∼7 x 10^6^ cells/mL), high-quality ROIs (i.e., ROIs containing a single in-focus particle) alone underestimated the total cell concentration as reported by flow cytometry, due largely to a significant increase in ROIs containing multiple objects compared to higher dilution levels. Accounting for the combined frequency of both multi-object ROIs and the estimated abundance of in-focus cells, a total abundance estimation that matched the quantity measured by flow cytometry was recovered. At low concentrations in this range (∼2 x 10^3^ cells/mL), replicate variability increased for ARTiMiS estimates as sample concentration approached the limit of quantification. The ARTiMiS’ limit of blank (LoB), determined from the false positive rate when processing blank samples, was 2.7 x 10^3^ particles/mL (**Text S3**). Testing was performed using flow cells with moderate prior use to simulate field-relevant conditions (e.g., fouling, debris adhesion, etc.). From the LoB and the procedure described in Text S3, the theoretical limit of detection was calculated to be 3.2 x 10^3^ particles/mL, though ARTiMiS yielded order-of-magnitude accurate estimates slightly below this threshold (**Figure 4)**. The limit of quantification was conservatively determined to be approximately 5 x 10^3^ particles/mL, based on the criteria described in Text S3.

### ARTiMiS Exhibited High Accuracy for Low-Complexity Microalgal Classification

“Off-the-shelf” convolutional deep neural network (CNN) models (ResNet^29^, MobileNet^30^, YOLO^31^, etc.), despite being publicly available, were designed to perform well on broad classification tasks often based on diverse, socially relevant targets (e.g., animals, vehicles, etc.). Identifying biological targets represents a narrower problem scope, and thus lower-complexity classification models^24,32^ have been developed for cell identification. Often, these architectures have lower memory and processing requirements; aiming to implement classification onboard ARTiMiS for real-time prediction, the instrument’s compute hardware constrained viable classifier complexity. The implemented “micro” CNN model architecture was modified from architectures described previously^24,32^, with alternating convolution and max-pooling layers (see **Materials and Methods**) and 62,000 total trainable parameters (for reference, MobileNet V2, designed to run in real time on smartphones, has 3 million trainable parameters).

Interspecies classification experiments were conducted using images obtained from *Scenedesmus obliquus* and *Chlorella sorokiniana* cultures (**Figure 5A**). While ellipsoidal *Scenedesmus* contrast with circular *Chlorella*, intra-class morphological diversity was high for both organisms (**Figure 5A - insert**), resulting in flattened feature distribution curves and long tails (**5A**). We therefore hypothesized that a random forest model might fail to accurately distinguish between the two classes and implemented a CNN classifier of the low-complexity architecture described. Against expectations, random forest achieved (**Figure 5B)** accuracy equivalent to or better than the CNN classifier (**Figure S8A**), achieving 98.5% and 97.9% accuracy on the test dataset, respectively. Ellipsoid-related features dominate the features of highest decision importance (**Figure 5C**), rather than size-related features as seen previously, which coincides with the visual intuition of archetypal class representations. The lack of a clear decision boundary between classes (compared to visible thresholds in **Figure 3B**) emphasized a key advantage of utilizing machine learning to identify patterns in N-dimensional feature spaces that are not visible in 2D or 3D visualizations, thus removing the burden of manual feature curation.

By creating pre-defined mixes of the two classes from sample data entirely independent from the training and test data, ranging in proportional ratio from 100:0 of one species to 0:100 of the other species with pre-set intervals in between, a model’s bias for one class or the other can be estimated. A small but consistent bias for the *Scenedesmus* class was observed for the CNN classifier (**Figure S8B**), and to a lesser extent the random forest classifier (**Figure S8C**). For example, in a sample containing 100% *Chlorella* cells, the CNN model erroneously estimates approximately 7% are *Scenedesmus*; at a 50:50 ratio, the classifier over-estimates *Scenedesmus* proportionally, by ∼3%. With equivalent accuracy, less bias, and lower computational resources required, random forest was shown to be a preferred method of taxonomic identification in this binary cell classification task. In quantifying the classifier’s bias, bias information can be accounted for in post-processing and corrected; performing this same correction for a human annotating live samples is considerably more challenging to quantify and consistently apply over time. Thus, a taxonomist’s input may only be required when curating training datasets; thereafter, a machine vision model can be trained, its biases quantified, and then a corrected model can be applied to predict the taxonomic composition of samples near-instantaneously. Training datasets can and should be updated routinely, though this incremental effort can be scaled to large and ongoing sample datasets.

### ARTiMiS Demonstrated Real-Time, Long-Term Classification Capabilities on Complex Microalgal Communities

To test ARTiMiS’ potential to monitor more complex microalgal communities, we used ARTiMiS to image samples from the EcoRecover wastewater nutrient recovery system, a full-scale microalgae-based nutrient removal process in the Village of Roberts, Wisconsin, USA^23^. Through a combination of direct observation and referencing of 18S rRNA gene sequencing^33^, the dominant taxonomic groups present in the EcoRecover microalgal community were identified and libraries comprising representative images of each group were annotated. Three clades of Chlorophyta were found to dominate the system at different times of the year: multiple members of the *Scenedesmaceae* family, one or several species of *Chlorella*, and two or more species of *Monoraphidium*. An additional noteworthy class of particles, primarily consisting of flocs of bacteria that either entered the system from the upstream secondary wastewater treatment process or developed in the EcoRecover process itself, were annotated. This “bacterial floc” class also included colonial cyanobacteria occasionally observed in the system in low abundances. A fifth null class, comprising out-of-focus objects or other particles of unknown identity, was created as a catch-all negative class (referred to as “Unknown”). As both the *Chlorellaceae* and *Scenedesmaceae* families were observed in high abundances in the EcoRecover mixed microalgal community, the previous binary classification technique was repeated using annotated samples from the EcoRecover dataset (**Figure S9A, S9B**) to determine if the conclusions from **Figure 5** would apply to this more complex community. Potentially owing to the greater morphological diversity observed in the field samples (**Figure S9C**), including the presence of multiple species within each group, the random forest binary classifier demonstrated significantly poorer accuracy (85%) as compared to the CNN classifier trained on the same training data (95%). This performance trend continued when including the other three classes in a multi-class classification task (**Figure S9D**). The intra-class morphological diversity combined with inter-class feature similarities likely made clean decision boundaries based on the measured semantic features infeasible and could be the reason for poor performance of the random forest classifier relative to the CNN model. The CNN, by contrast, was trained to develop its own “features” to distinguish between the classes by virtue of the deep learning technique.

Higher accuracy was achievable with a CNN classifier, which yielded per-class accuracy greater than 90% for each of the target classes (**Figure 6A**). While lack of transparency regarding the “features” used for classification represents a challenge for model interpretability and troubleshooting, the direct input data (**Figure 6B**) for a CNN classifier remains visually intuitive to a human data curator, helping to partially intuit what the classifier “sees.” After training, the classifier was applied to continuous sampling periods to construct a time-series view of the microalgal community (**Figure 6C**). Views of the four distinct time series capture mixed microalgal community dynamics only observable with daily sampling resolution. These patterns include a stratified community with stable total biomass (January 15 - February 11, 2022), a stratified community with oscillating total biomass (October 5-31, 2022), succession of one dominant taxonomic group to another during a period of stable total biomass (April 21 – June 2022), and a total biomass system recovery dominated by a single taxonomic group (July 25 – September 8, 2022). These dynamics, both of total biomass and of community composition, would traditionally necessitate measurement by two separate analytical techniques: biomass characterization via total suspended solids, optical density, fluorometry, etc., and DNA sequencing analysis for taxonomy. The ARTiMiS, however, enabled both to be measured by a single instrument. Using ARTiMiS, the only operator inputs required were sample collection, loading of the instrument, and running a pre-set sample processing configuration, a workflow requiring 3-5 minutes of operator time in total. Furthermore, ARTiMiS provided an archive of image data available for reference months after sample collection, to provide a comparison between sample appearance or re-analysis.

**Figure 6:**
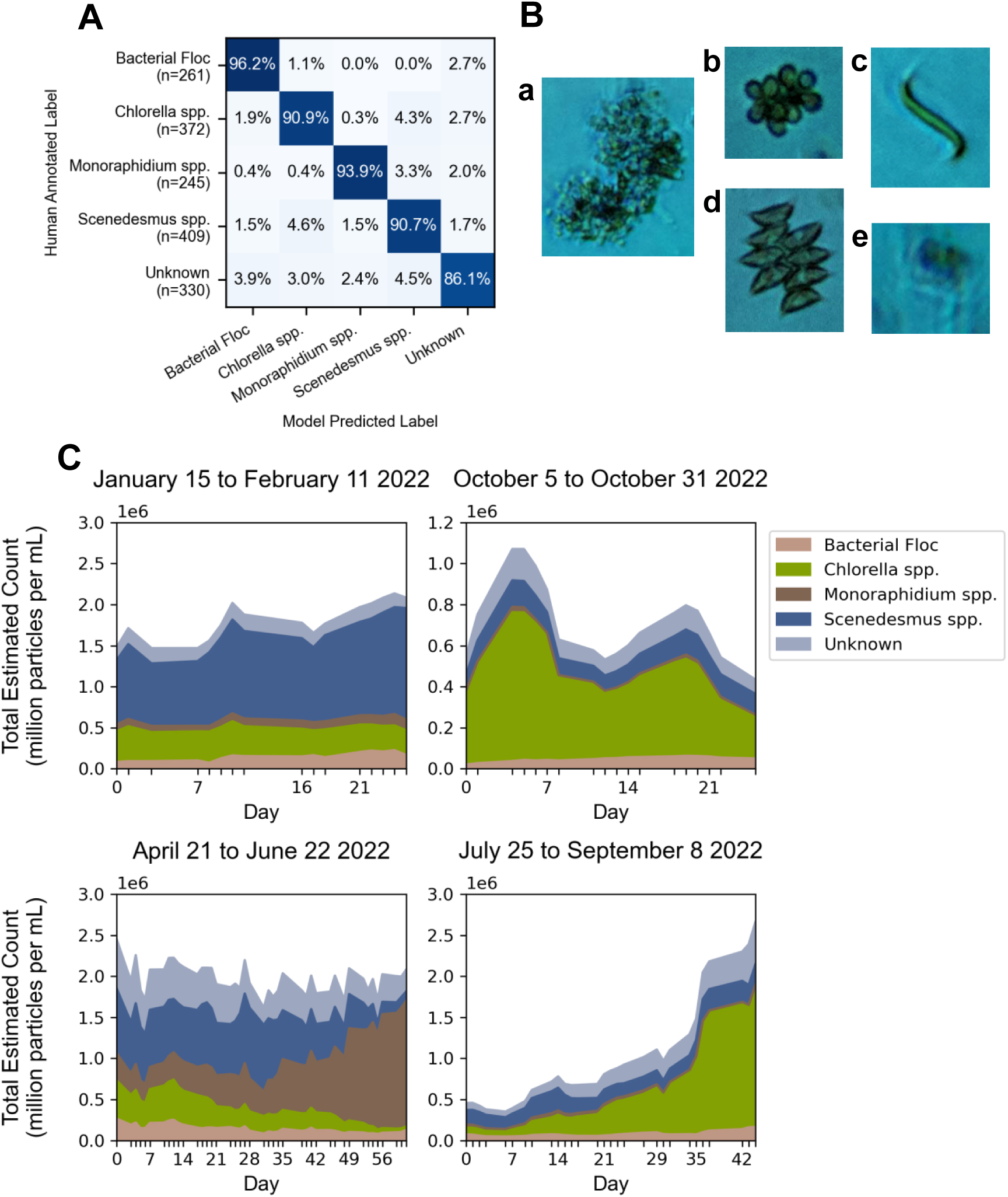
ARTiMiS used to process samples of a mixed microalgal community. A Convolutional Neural Network (CNN) classification model was trained to distinguish between four dominant taxonomic groups and one null class, “Unknown,” (A) and achieved 90% accuracy or greater for each of the target classes (and 86.1% for the “Unknown” class) when evaluated on unseen test data. Representative images of each class of organisms are shown (B): Bacterial Floc (a); Chlorella spp. (b); Monoraphidium spp. (c); Scenedesmus spp. (d); Unknown (e). Predicting taxonomic composition of time-series collected samples (C) across different seasonal periods allowed for observation of a variety of behaviors: stable biomass, stratified community (upper left); variable biomass, stratified community (upper right); stable biomass, dynamic community (lower left); growing biomass, stratified community (lower right).

### The ARTiMiS is a Low-Cost, Real-Time Flow Imaging Microscopy Solution with Low Technical Barriers to Adoption

Data from semi- or fully-automated digital imaging microscopy instruments has been consistently mined for machine learning proof-of-concept studies in the last two decades. Yet, manual microscopy remains a mainstay in most phycological research laboratories and industrial-scale microalgal cultivation operations^34,35^. The rationale for incomplete adoption of automation varies for each end user or organization, but can generally be summarized as one or several of the following constraints: (1) workflows that require a deeper level of microscopist-sample interaction, (2) information barriers to options available, (3) financial barriers to adopting expensive commercial solutions, and/or (4) technical barriers to adoption of open source and open hardware solutions that require both engineering and phycology-related skillsets to build, use, and maintain them. While the ARTiMiS was developed contemporaneously with other digital microscopy platforms and other phycology-focused machine learning workflows^36–39^, its design motive was distinct. There remains a gap among options available to users who would benefit from automated digital microscopy: turnkey solutions require significant financial investment, and even still may lack modern features such as built-in machine learning tools to automate identification. The design impetus of ARTiMiS was to fill this gap: to provide a solution that was financially accessible but did not as a caveat expect a user to refer to its source code and design drawings to literally build the solution on their own. The current study systematically demonstrates ARTiMiS’ applications from low complexity laboratory to high complexity industrial microalgal systems; with appropriate adaptations, the ARTiMiS can be extended to other environments where real-time phytoplankton monitoring is necessary and desired (e.g., harmful algal blooms, etc.).

## Acknowledgements.

This research was supported by the National Science Foundation (NSF) Graduate Research Fellowship to B.G. and by the Paul L. Busch Award to A.P. This research was also supported by NSF (CBET Award Number: 2220792) and by the U.S. Department of Energy, Office of Energy Efficiency and Renewable Energy (Award Number: DE-EE0009270).

