## Supplemental materials for "Introducing ARTiMiS: A low-cost flow imaging microscope for phytoplankton monitoring in engineered and natural environments"

**Supplementary Text S1: Fluidic, optical, actuation, and automation systems design and construction.**

The device housing was modeled with SolidWorks (Dassault Systemes) to be 3D printed on a Raise3D E2 3D printer (Raise3D Technologies, Inc.) using polylactic acid (PLA) filament for rigid components and polyethylene terephthalate glycol (PETG) for components benefitting from added flexibility. Power is supplied by a 12V 3A DC power supply in barrel jack format, stepped down to 5V for appropriate components with a DC-DC converter. To drive motors and actuators, off-the-shelf motor driver control boards are used for stepper motor control and for pulse width modulation (PWM) motor control. To bind the microcontroller, actuator, and power systems, a printed circuit board (PCB) for connection routing was designed using the design software EasyEDA (LCSC Electronics) and fabricated with a combination of surface mount and through hole technology elements.

The core element of the fluidic system was a commercially available microfluidic chip. The COC-polymer microfluidic chip (Fluidic 142, Microfluidic ChipShop) contains 16 parallel channels of dimensions 1000 μm (width) x 200 μm (depth) x 18 mm (length) with extruded interfaces to attach 0.76 mm inner diameter silicone tubing. The device's internal liquid volume measures approximately 450 μL, thus sample volumes as small as 0.5 mL can be processed via recirculation. The enclosed fluid imaging volume, measured as the extents of the optical field of view through the full microfluidic channel depth, is 0.060 μL. For continuous flow, microfluidic tubing attached at the inlet and outlet of the channel, which was gated by electronic pinch valves on both sides. The instrument was designed to interface with a sample source with positive pressure from inlet to outlet, which could, for example, be supplied by gravity. For deployment scenarios without positive pressure, such as when the instrument is configured for benchtop operation, a low-cost peristaltic pump was used to pull samples through the fluidic system. The 30 µm depth of field, while sufficient to capture in-focus images of planar objects, was much narrower than the full imaging volume depth (200 µm). As in cell counting chambers (e.g., Sedgewick Rafter Cell), particle settling was used to maximize the number of objects in focus within a given image frame.

Pinch valves were added to hold the image volume stationary to allow particles to settle in the microfluidic chip prior to imaging. Settling time was variable based on sample material and particle density, though typically ranged between 30-90 seconds and is configurable by the instrument user for each sample run.

The optical system was built around a reversed lens configuration of two M12-threaded board lenses. These lenses are often employed in closed-circuit security systems, DIY electronics projects, and contemporary low-cost microscopy projects (Li et al., 2019; Pollina et al., 2022) due to their small form factor and extremely low cost. A short focal length (2.8 mm) lens was selected as the tube lens, and a longer focal length (16 mm or 25 mm) lens was selected as the objective in this system. These lens configurations provide theoretical magnification of 5.7X and 8.9X, respectively, calculated as the focal length ratio. Empirical assessment of magnification indicated that effective magnification was slightly lower, approximately 5X and 8X, respectively (**Figure S1**).

The lenses are mounted via a commercially available M12 thread adapter to an 8MP Raspberry Pi Camera v2 (Sony IMX219 sensor, stock lens removed), with a sensor-side pixel width of 1.12 µm. Illumination is provided by the off-the-shelf UnicornHAT HD (Pimoroni® PIM273) 16x16 programmable LED array. Individually addressable red, green, and blue diodes comprised each LED, programmatically supported by a publicly available Python library (unicornhathd 0.0.4, PyPI) to drive the LED panel. Use of this light source enables serial imaging of the same sample volume under bright field, dark field, and other illumination conditions without mechanical movement, thus reducing complexity and improving system reliability. One impact of using non-traditional optical elements was the introduction of visual artifacts, which required post-processing correction (**Figure S2**). One of the notable artifact was the inconsistent color field of bright field images: a blue-shifted image with a red vignette near the periphery. This coloration was the combined result of using an RGB LED light source, rather than a pre-tuned white LED, in conjunction with the Raspberry Pi Camera V2's color correction firmware. As a result of removing the camera's stock lens in exchange for board lenses to create the desired magnification for microscopy, the camera's native lens shading firmware over-corrects the resulting image, as observed elsewhere in (Bowman et al., 2020; Pollina et al., 2022). Bowman and colleagues study this effect in detail and offer both a comprehensive description of the problem and present a number of solutions. In order to produce post-processed images with consistent illumination and color, a combined flat-field and color correction step is applied. A post-capture processing approach was selected in order to enable retroactively applying the correction in a consistent manner across all datasets, and to enable updates to the correction algorithm to improve results without permanently modifying the underlying raw image data. The flat field and color correction method used here operates as follows: first, a collection of "blank" bright (***B***) and dark (***D***) field reference images are collected at a range of illuminator brightness intensity levels (ranging from 40% to 100% maximum brightness in steps of 10%) to provide reference images at a number of different operational illumination settings. To correct a new bright field image, a reference of equivalent illumination intensity is selected. The average pixel intensity across each color channel is measured from the reference bright and dark field images ($\bar{B}, \bar{D}$), and combined brightness and color gain is calculated as the difference of color channel means over the per-pixel image differences, multiplied by a color gain parameter. In most cases, a color gain vector of [0.5, 1, 1] is applied to the R, G, and B channels, respectively. The corrected image ($\hat{\boldsymbol{I}}$) is calculated as the difference between the raw image ($\boldsymbol{I}$) and the dark field reference image ($\boldsymbol{D}$) multiplied by the per-channel gain vector ($\vec{G}$). An example before and after corrected image is shown in (**Figure S2**). As a final step, a small watermark is written to one corner of the image for provenance tracking, to denote that the image has been corrected when saved to disk in order to prevent correcting it again in automated workflows.

$$\vec{G} = \frac{\bar{B}-\bar{D}}{\boldsymbol{B}-\boldsymbol{D}} \cdot \vec{C} \ (gain)$$

$$\hat{\boldsymbol{I}} = (\boldsymbol{I} - \boldsymbol{D}) \cdot \vec{G} \ (image\ correction)$$

To enable remote and autonomous operation of the imaging system, a stepper motor linear actuator inspired by the Octopi project (Li et al., 2019) was used to govern focusing (21H4U-2.5-907, Haydon Kerk Pittman, AMETEK Inc.). Automated fluid handling was controlled via a low-cost, 2 mm inner diameter peristaltic pump (AE1207, Gikfun), and two normally closed electronic pinch valves (NPV2-1C-02-12, Clippard). A stop-flow pattern was implemented to minimize particle motion within the field of view during exposure time-constrained image captures. Both motor controllers were supplied power and connected to the Raspberry Pi's general purpose input/output (GPIO) pins via a custom-designed printed circuit board to route power and control signals between the electronic elements. Drivers for each motor controller were created in Python to enable both automated and manual control. For a complete list of commercially available hardware components used, see **Table S1**.

**Supplementary Text S2: Image pre-processing for object detection.**

RGB color images were converted to grayscale and minimum-maximum normalized to the range [0,1]. A Difference of Gaussians filter was applied prior to foreground-background thresholding by a fixed value. After binarizing the image as foreground-background, a series of dilation, small hole removal, and erosion steps were applied to solidify pixel regions corresponding to a single object using OpenCV (opencv-python, v4.6.0). Contiguous foreground pixel regions were labeled as separate objects (scikit-image, v0.19.2), which were subsequently screened by size to remove artifacts such as edges of the flow cell if visible in the frame. Images are converted to grayscale and min-max normalized, a less computationally expensive noise reduction step is applied where two white top-hat transforms are used with different structuring elements and subtracted to create a denoised image. The algorithm is provided with a target region of interst (ROI) size (e.g., 100 pixels) and identifies local luminance intensity maxima of distances proportional to that tile size. Luminance intensity peaks are filtered by a value threshold to remove dim local maxima, and peaks adjacent to the edge of the frame are discarded. These coordinates are used to establish preliminary ROIs of pre-set size centered on the luminance intensity peak, and are adjusted to the optical center of mass from the un-filtered image frame. As ROI coordinates frequently shift with this adjustment, overlapping ROIs are merged to minimize double-counting the same object.

**Supplementary Text S3: Limit of blank, detection, and quantitation.**

To establish LoB, seven blank samples (Milli-Q filtered water, Millipore Sigma) were processed on ARTiMiS for a total of 20 image frames each. Object density was estimated using the Fast Object Detection algorithm. LoB was calculated according to the equation described by Armbruster and Pry:

$$\text{LoB} = \text{mean}_{\text{blank}} + 1.645(\text{SD}_{\text{blank}})$$

To establish LoD and LoQ, a serial dilution of laboratory-cultivated *Chlorella sorokiniana* (UTEX1602) was performed with parabolic dilution series (12x-46080x). As described by Hubaux and Vos (1970) this methodology increases sample point density near the upper and lower limits of quantitation to efficiently capture more information in the non-linear calibration regions. Due to the time requirement of sample processing on ARTiMiS, sample points were diluted in nutrient-free Bold's 1NV medium, and cultures were sampled near stationary phase, to mitigate impacts of cell division on results. The number of widefield frames required for each dilution step was estimated using hemocytometer sampling protocols (Patterson, 1979) targeting a minimum of ~100 cells per sample point. At higher concentrations where 100 or more cells were expected in a single frame, a minimum of five frames were captured. Limit of Quantitation definitions are dependent on constraints for bias and reproducibility between replicates. Here, we define LoQ as the minimum sample concentration where the sample mean measured by an external gold-standard method (i.e., flow cytometery) is within a 95% confidence interval established by an ARTiMiS-estimated sample concentration, and a coefficient of variation (CV) ≤ 20%. Theoretical Limit of Detection was estimated according to the subsequent equation:

$$\text{LoD} = \text{LoB} + 1.645(\text{SD}_{\text{low concentration sample}})$$

### Supplementary Tables

**Table S1:** Hardware Bill of Materials (BOM) for assembling a complete ARTiMiS instrument, including peripheral display for hardwired display output.

| Hardware Bill of Materials | | | | | | |
|---|---|---|---|---|---|---|
| **BOM No.** | **Name** | **Qty.** | **Line Item Cost** | **Description** | **Vendor Product No.** | **Link** |
| 1 | M2 Heatset Insert | 8 | $ 1.41 | | 94459A110 | https://www.mcmaster.com/94459A110 |
| 2 | M2.5 Heatset Insert | 8 | $ 0.90 | | 94180A321 | https://www.mcmaster.com/94180A321 |
| 3 | M3 Heatset Insert | 13 | $ 2.49 | | 94459A130 | https://www.mcmaster.com/94459A130 |
| 4 | 1/16 x 3/16 Neodymium Magnet | 30 | $ 2.79 | | 5862K139 | https://www.mcmaster.com/5862K139 |
| 5 | M2.5 11mm 6mm brass standoff | 4 | $ 1.17 | | | https://www.amazon.com/gp/product/B07KM27KC6/ |
| 6 | M2.5 5mm 5mm brass standoff | 4 | $ 1.40 | | | https://www.amazon.com/40Pcs-Project-Brass-Standoff-Spacers/dp/B073X9FZZT |
| 7 | M2 6mm machine screw | 6 | $ 0.32 | M2-0.40mm Machine Screw, Cheese, Phillips, A4 Stainless Steel, Plain, 6mm Length | | https://www.grainger.com/product/FABORY-M2-0-40mm-Machine-Screw-31JR95? |
| 8 | M2 14mm machine screw | 2 | $ 0.09 | M2-0.40mm Machine Screw, Cross Recessed Raised Cheese, Phillips, A2 Stainless Steel, Plain, 14mm Length | 38EA23 | https://www.grainger.com/product/FABORY-M2-0-40mm-Machine-Screw-38EA23 |
| 9 | M2.5 5mm screw | 8 | $ 0.40 | | | https://www.amazon.com/40Pcs-Project-Brass-Standoff-Spacers/dp/B073X9FZZT |
| 10 | M3 Pan head 6mm screw | 12 | $ 0.72 | | 90604A722 | https://www.mcmaster.com/90604A722/ |
| 11 | M3 Thumbscrew | 1 | $ 2.37 | | 92552A414 | https://www.mcmaster.com/92552A414/ |

| | | | | | | |
|---|---|---|---|---|---|---|
| 12 | M3 Lock washer | 2 | $ 0.04 | | | https://www.amazon.com/uxcell-500pcs-Stainless-Spring-Washer/dp/B018TG7DNS/ |
| 13 | #4-40 3/16" machine screws | 4 | $ 0.11 | | 1ZB34 | https://www.grainger.com/product/FABORY-4-40-Machine-Screw-1ZB34 |
| 14 | M12 lens mount adapter | 1 | $ 8.50 | M12 lens - Pi Camera V2 adapter mount | PT-LH024RPM | http://www.m12lenses.com/CNC-Machined-Raspberry-Pi-M12-Lens-Holder-Metal-p/pt-lh024rpm.htm |
| 16 | 4mm ID Hose Clamps | 2 | $ 1.30 | uxcell 4mm Inner Dia Spring Clip Water Pipe Fuel Line Hose Clamps | | https://www.amazon.com/gp/product/B06XKJGTTX/ |
| 17 | Mini-Luer Male Fluid Connector Blue | 2 | $ 5.13 | | 10000096 | https://labsmith.com/products/male-mini-luer-fluid-connector/?sku=10000096 |
| 18 | Silicone Tubing 0.76mm ID / 1.65mm OD | 0.2 | $ 2.60 | | 10000031 | https://labsmith.com/products/silicone-tubing-0-76-mm-id-p-n-10000031/ |
| 19 | Silicone Tubing 2mm ID / 4mm OD | 0.1 | $ 0.50 | | | https://www.amazon.com/gp/product/B0852HZSTR/ |
| 20 | Peristaltic Pump | 1 | $ 12.98 | 12V DC Dosing Pump | | https://www.amazon.com/Gikfun-dosificadora-conector-Laboratorio-Anal%C3%ADtico/dp/B01IUVHB8E/ |
| 21 | 16-Channel Microfluidic Chip | 1 | $ 48.87 | straight-16-channel-olive-chip | 10000278 | https://products.labsmith.com/straight-16-channel-olive-chip-p-n-10000278/ |
| 22 | Solenoid Pinch Valve | 2 | $ 94.02 | 2-Way N.C. Pinch Valve, 0.75" Dia., 0.030" ID-0.065" OD Tubing, 12 VDC | NPV1-1C-01-12 | https://www.clippard.com/part/NPV1-1C-01-12 |
| 24 | HKP Nema 8 Captive Linear Actuator | 1 | $210.19 | Captive linear actuator for focusing | 21H4U-2.5-907 | https://prototypes.haydonkerk.com/ecatalog/hybrid-linear-actuators/en/linear-actuator-21H4U-2.5-907 |
| 25 | Raspberry Pi Camera Module V2 | 1 | $ 27.45 | 8MP R Pi Camera | | https://www.amazon.com/Raspberry-Pi-Camera-Module-Megapixel/dp/B01ER2SKFS |
| 26 | Pimoroni Unicornhat HD | 1 | $ 34.95 | Programmable LED array | 3580 | https://www.adafruit.com/product/3580 |
| 27 | M12 2.8mm board lens | 1 | $ 6.25 | M12 2.8mm board lens | PT-02820 | http://www.m12lenses.com/2-8mm-F2-0-Board-Lens-p/pt-02820.htm |

| | | | | | | |
|---|---|---|---|---|---|---|
| 28 | M12 16mm board lens | 1 | $ 6.50 | M12 16mm board lens | PT-1620 | http://www.m12lenses.com/16mm-F2-0-Board-Lens-p/pt-1620.htm |
| 29 | M12 25mm board lens | 1 | $ 15.50 | Focal Length 25mm, F2.0, M12*0.5 Board Lens | PT-2520 | http://www.m12lenses.com/25mm-F2-0-Board-Lens-p/pt-2520.htm |
| 31 | Raspberry Pi 4 B 4GB | 1 | $ 55.00 | Single board computer | 2GB-9003 | https://www.pishop.us/product/raspberry-pi-4-model-b-2gb/ |
| 32 | 32GB Sandisk microSD card | 1 | $ 10.29 | Onboard storage device | | https://www.amazon.com/dp/B06XWMQ81P/ |
| 34 | LM2596 DC-DC Buck Converter Step Down Module | 2 | $ 3.60 | Step 12V to 5V | | https://www.amazon.com/Zixtec-LM2596-Converter-Module-1-25V-30V/dp/B07VVXF7YX/ |
| 35 | 12VDC 5.5mm x 2.1mm barrel jack | 1 | $ 0.37 | Barrel jack to screw terminal for board interface | | https://www.amazon.com/Power-Connector-Female-Adapter-Camera/dp/B07C61434H/ |
| 36 | 16mm latching power button | 1 | $ 3.00 | | | https://www.amazon.com/gp/product/B08MPZSDWL/ |
| 37 | 12VDC 3A power supply | 1 | $ 9.99 | | | https://www.amazon.com/gp/product/B07VQGHSWY/ |
| 39 | Acuator controller PCB | 1 | $ 1.89 | | | https://jlcpcb.com/ |
| 40 | Allegro A4988 bipolar motor stepper | 1 | $ 1.80 | Stepper motor driver for bipolar linear actuator | | https://www.amazon.com/BIQU-Compatible-StepStick-Stepper-Controller/dp/B01FFFYVV8/ |
| 41 | TB6612FNG Dual Motor Driver Carrier | 1 | $ 3.13 | Motor driver for DC motors, valves | 713 | https://www.pololu.com/product/713 |
| 42 | Female 2.54mm pitch pin headers, 1x8 | 4 | $ 0.30 | | | https://www.amazon.com/gp/product/B07PLBC2GT |
| 43 | Male 2.54mm pitch pin headers (variable) | 1 | $ 0.02 | | | https://www.amazon.com/Jabinco-Breakable-Header-Connector-Arduino/dp/B0817JG3XN/ |
| 44 | Male Right Angle 2.54mm 2x20 pin headers | 1 | $ 0.37 | | | https://www.amazon.com/gp/product/B07VSC4PWW/ |
| 45 | 2 pin screw terminals | 2 | $ 0.70 | | | https://www.amazon.com/DIYhz-green-Terminal-Connector-Arduino/dp/B0774YRVVX/ |
| 46 | Female-Female Jumper Wires, 15cm, Pack | 1 | $ 2.50 | | | https://www.amazon.com/gp/product/B07GCZVCGS/ |

| | | | | | | |
|---|---|---|---|---|---|---|
| 48 | PLA 1.75mm 3D Printing Filament (per kg) | 0.4 | $ 12.25 | | | https://www.raise3d.com/products/r3d-premium-pla-filament/ |
| 50 | microHDMI to HDMI cable | 1 | $ 5.99 | | | https://www.amazon.com/Adapter-Wenter-Action-Camera-Supported/dp/B07VRCK5W1/ |
| 51 | 10.5 inch portable FHD Display | 1 | $ 69.42 | | | https://www.amazon.com/10-5inch-Portable-1920x1280P-External-Ultra-Thin/dp/B0C7KPPXTZ/ |
| | **Total Cost** | | **$669.54** | | | |

**Table S2:** Features measured during semantic feature extraction. Features that are derived entirely or predominantly from an external computer vision library are indicated by dependency source. Features without a cited library dependency are derived using mathematical equations or methods from the Python standard library.

| Feature Name | Description | Library Dependency |
|---|---|---|
| UUID | Universally Unique Identifier, unique for each particle. | standard |
| XY Coordinates | Image coordinates of center of crop region. | standard |
| Area | Sum of count of pixels comprising dominant object in foreground. | scikit-image |
| Bounding Box Area | Area of the smallest rectangle enclosing the dominant object. | scikit-image |
| Convex Area | Area of the convex hull of the dominant object. | scikit-image |
| Total Area | Sum of all pixels in binary image foreground (can include other objects). | standard |
| Patch Size | Size of the image crop (ROI), in pixels and microns. | standard |
| Mean Intensity | Mean pixel brightness value across entire ROI. | scikit-image |
| Maximum Intensity | Maximum pixel brightness value across entire ROI. | standard |
| Object Intensity Sum | Sum of pixel brightness values for all pixels comprising the dominant object. | standard |
| Total Intensity Sum | Sum of all pixel brightness values across the entire ROI. | standard |
| Solidity | Ratio of pixels in the enclosing region to pixels of the convex hull image. | scikit-image |
| Perimeter | Perimeter contour calculated through the centers of border pixels using a 4-connectivity. | scikit-image |
| Equivalent Diameter | Diameter of a circle with the same area as the region. | scikit-image |
| Feret Diameter (max) | Maximum Feret's diameter, the longest distance between points around the convex hull. | scikit-image |
| Major Axis Length | Length of major axis of an ellipse having the same normalized second central moments as the dominant object. | scikit-image |
| Minor Axis Length | Length of minor axis of an ellipse having the same normalized second central moments as the dominant object. | scikit-image |
| Eccentricity | Ratio of the focal distance over the major axis length, calculated from an ellipse having the same normalized second central moments as the dominant object. | scikit-image |
| Orientation | Angle between the 0th axis and major axis of the ellipse having the same normalized second central moments as the dominant object. | scikit-image |
| Centroid | Weighted centroid of the image. | scikit-image |
| Hu Moments | Image moments: translation, scale, and rotation invariant. | scikit-image |

| | | |
|---|---|---|
| Hu Circularity | Circularity of object using the 0th Hu's moment as the object's radius. | scikit-image |
| Entropy | Shannon's entropy of the image. | scikit-image |
| Circularity | Circularity of object using the object's calculated area and perimeter. | standard |
| Euler Number | Euler characteristic of binary image. | scikit-image |
| Object Topography | Distance transform of object, binned as topographic levels. | scipy |
| Aspect Ratio | Ratio of minor axis length to major axis length. | standard |
| Biovolume (Sphere) | Calculated volume of object, assuming sphere (circular) geometry. | standard |
| Biovolume (Spheroid) | Calculated volume of object, assuming spheroid (elliptical) geometry. | standard |
| Edge Noise (Laplacian) | Blurriness of object as calculated as the standard deviation of Laplacian of Gaussian blur. | OpenCV |
| Edge Noise (Gradient) | Blurriness of object as calculated as the standard deviation of Sobel gradient of Gaussian blur. | OpenCV |
| Edge Gradient | Visual gradient at edge of object, calculated as average pixel intensity of object's outer edge. | standard |
| Edge Difference | Visual gradient at edge of object, calculated as difference in pixel values outside and inside the object's edge. | standard |

**Table S3**: Recall and processing times for Fast Detector and Segmenter algorithms, subdivided into high/low concentration Chlorella and EcoRecover. Recall refers to the same metric as “perfect crops” in Table 1: number of correctly identified regions of interest divided by total number of valid objects as annotated by a human observer.

Processing time values are reported as the mean of 6 replicated runs, (minimum time observed, maximum time observed) in seconds.

| | | Recall | Mean (Range) Time (PC) | Events per Second (PC) | Mean (Range) Time (Raspberry Pi 4) | Events per Second (Raspberry Pi 4) |
|---|---|---|---|---|---|---|
| Mono-culture [low] | Fast Detector | **97.3%** | **0.75s** (0.74s, 0.76s) | 104.0 | **6.22s** (6.18s, 6.27s) | 12.54 |
| | Segmenter | **94.6%** | **2.43s** (2.41s, 2.44s) | 25.51 | **35.62s** (35.58s, 35.67s) | 1.741 |
| Mono-culture [high] | Fast Detector | **84.3%** | **0.82s** (0.81s, 0.82s) | 340.2 | **6.29s** (6.27s, 6.33s) | 44.36 |
| | Segmenter | **84.6%** | **2.66s** (2.65s, 2.68s) | 108.6 | **35.54s** (35.48s, 35.61s) | 8.132 |
| Mixed Community [low] | Fast Detector | **66.2%** | **0.81s** (0.80s, 0.81s) | 64.20 | **5.83s** (5.81s, 5.87s) | 8.919 |
| | Segmenter | **92.9%** | **2.47s** (2.46s, 2.49s) | 14.57 | **38.21s** (38.01s, 38.31s) | 0.942 |
| Mixed Community [high] | Fast Detector | **69.9%** | **0.82s** (0.82s, 0.83s) | 211.0 | **6.06s** (6.05s, 6.08s) | 28.55 |
| | Segmenter | **95.3%** | **2.61s** (2.58s, 2.65s) | 49.43 | **36.28s** (36.18s, 36.36s) | 3.556 |

### Supplementary Figures

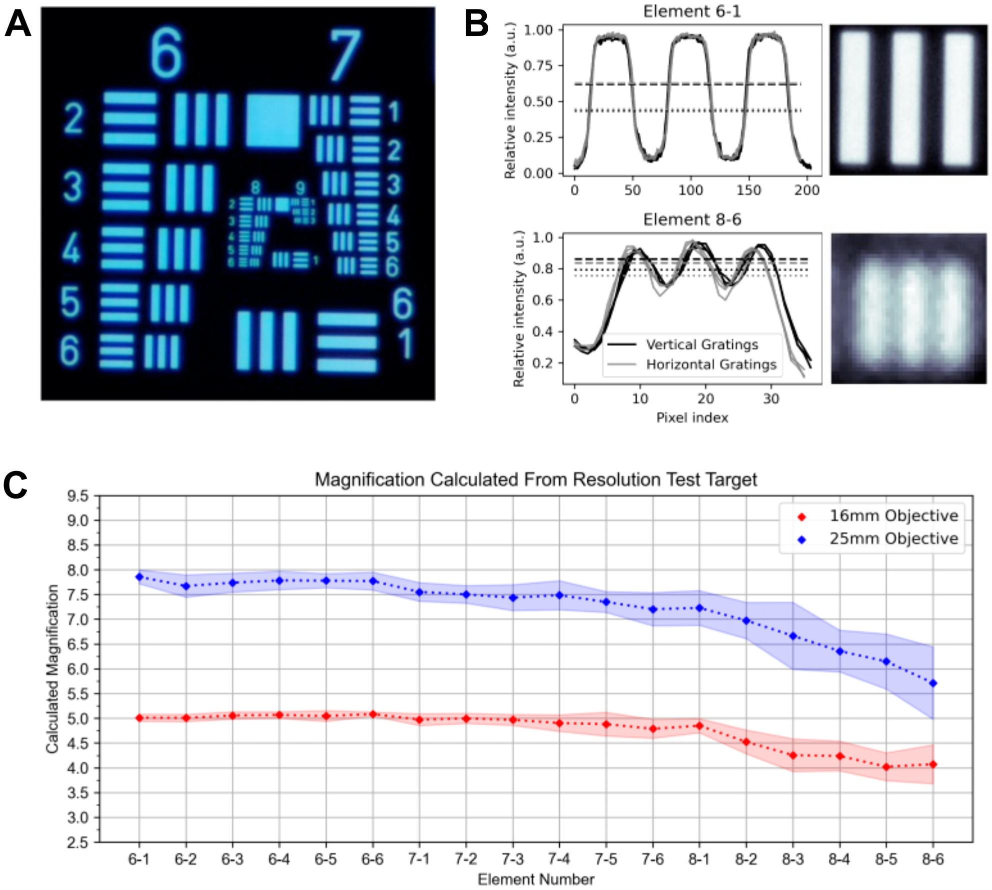


**Figure S1: (A)** USAF Resolution Test Target, bright field illumination. Title numbers describe Group, row numbers describe element number. **(B)** Pixel-wise measurement of line gratings to determine sensor-side line width, demonstrating grating silhouette for Group 6 Element 1 and Group 8 Element 6. Width measured as 60% of peak/trough magnitude, depicted by dashed and dotted lines. Black traces, vertical line pairs; gray traces, horizontal line pairs. **(C)** Magnification values calculated as the multiple of known line width to pixel width in resulting image (e.g., 5X describes a 10 μm line width occupying 45 1.12 μm pixels (50 μm total) width in the sensor-side image). Calculation was performed for each element in Groups 6-8 under 16 mm objective (red) and 25 mm objective (blue). Error envelopes represent standard deviation of mean.

### Flat Field and Color Correction

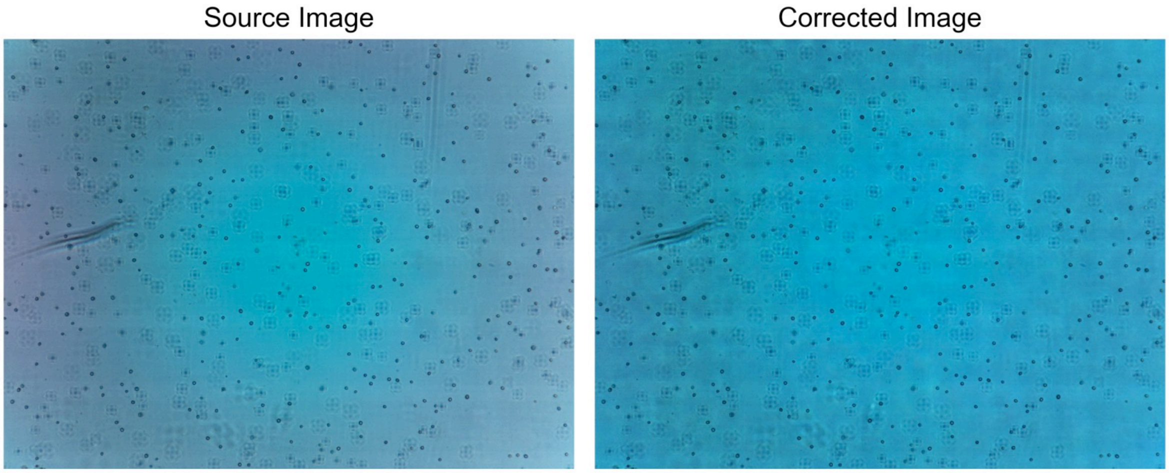


**Figure S2**: Representative depiction of before (left) and after (right) views when applying flat field and color correction to bright field images. Note color and luminance balancing of the dimmer red-tinted ring and lighter green-tinted center to a more uniform blue-tinted final image.

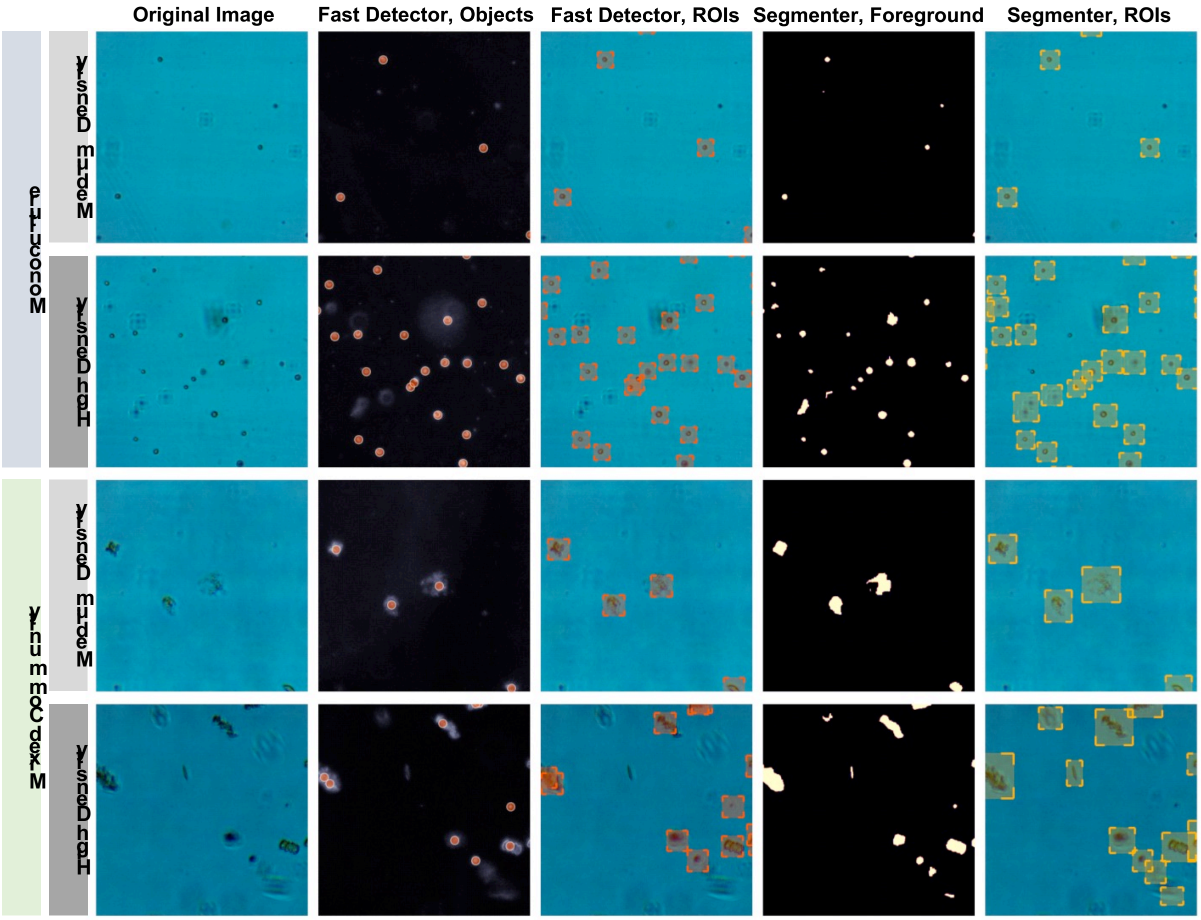


**Figure S3:** Object detection algorithm workflow representative examples. Rows pertain to different sample types, depicting both medium- and high-density samples for monospecific *Chlorella sorokiniana* culture and mixed microalgal community samples. Columns depict three key stages of each object detection algorithm: the original image, followed by the detected objects' coordinates (orange circle) and corresponding regions of interest, or ROIs, (orange rectangles) for the Fast Detector algorithm; the segmentation foreground (yellow mask) and corresponding ROIs (yellow rectangles) for the Segmenter algorithm.

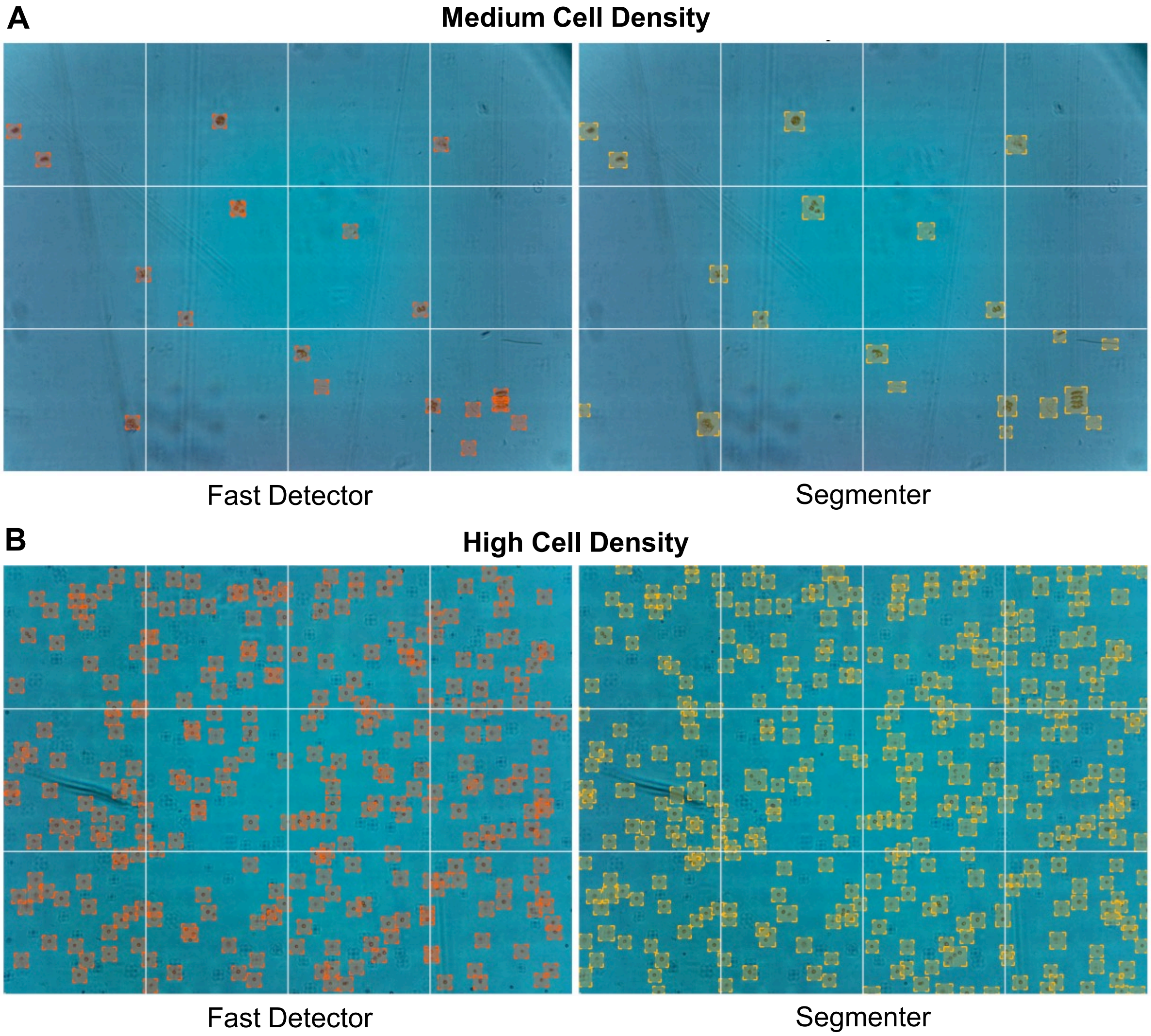


**Figure S4: (A)** Representative widefield frame of medium cell density sample, depicting regions of interest (ROIs) from Fast Detector (left) and Segmenter (right) overlaid on the same frame. White gridlines were drawn in post-processing to create a digital simulation of the hemocytometry cell counting technique. **(B)** Same layout as (A), depicting a high cell density sample.

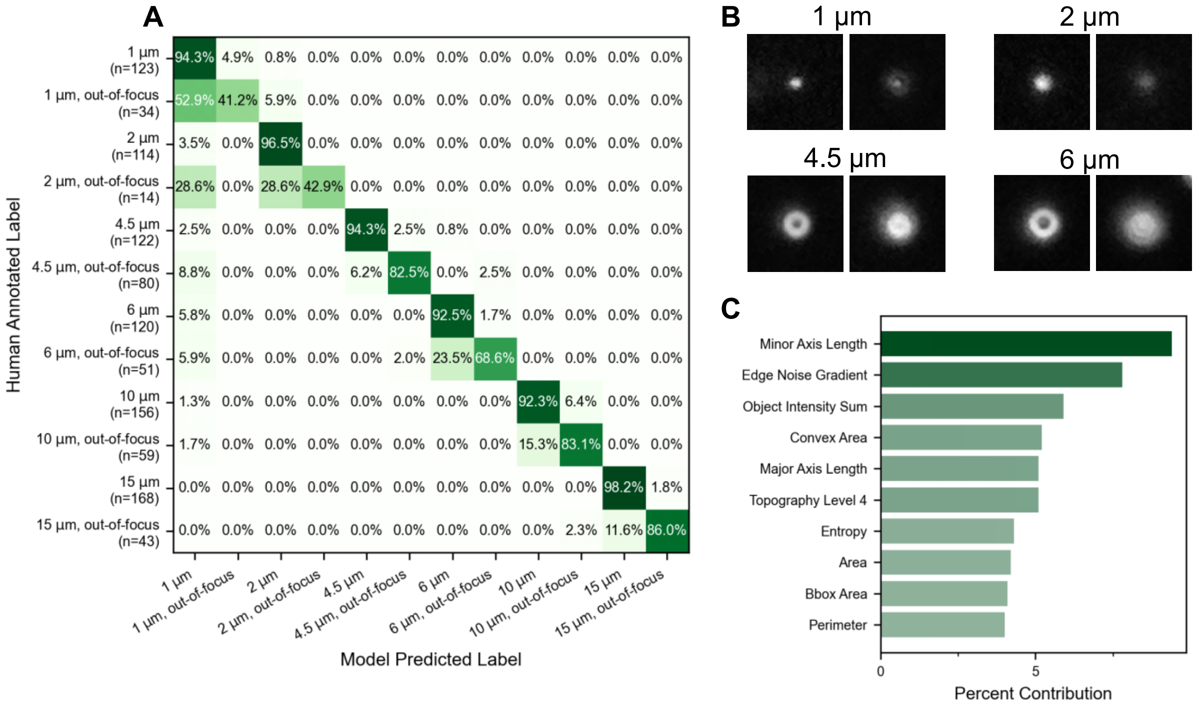


**Figure S5: (A)** Confusion Matrix describing random forest classification model performance when trained and tested on a dataset comprising both in- and out-of-focus calibration beads ranging in size from 1 to 15 µm. On-diagonal values describe correct model predictions, off-diagonal values represent misclassification rate on a per-class basis (e.g., 28.6% of 2 µm, out-of-focus particles are misclassified as 1 µm objects by the model). **(B)** Representative images of in-focus (left) and out-of-focus (right) examples of four of the particle size classes evaluated in (A). **(C)** Ranked importance of features for random forest classification. Dominant features for class differentiation overall describe particle size and degree-of-focus.

A

B

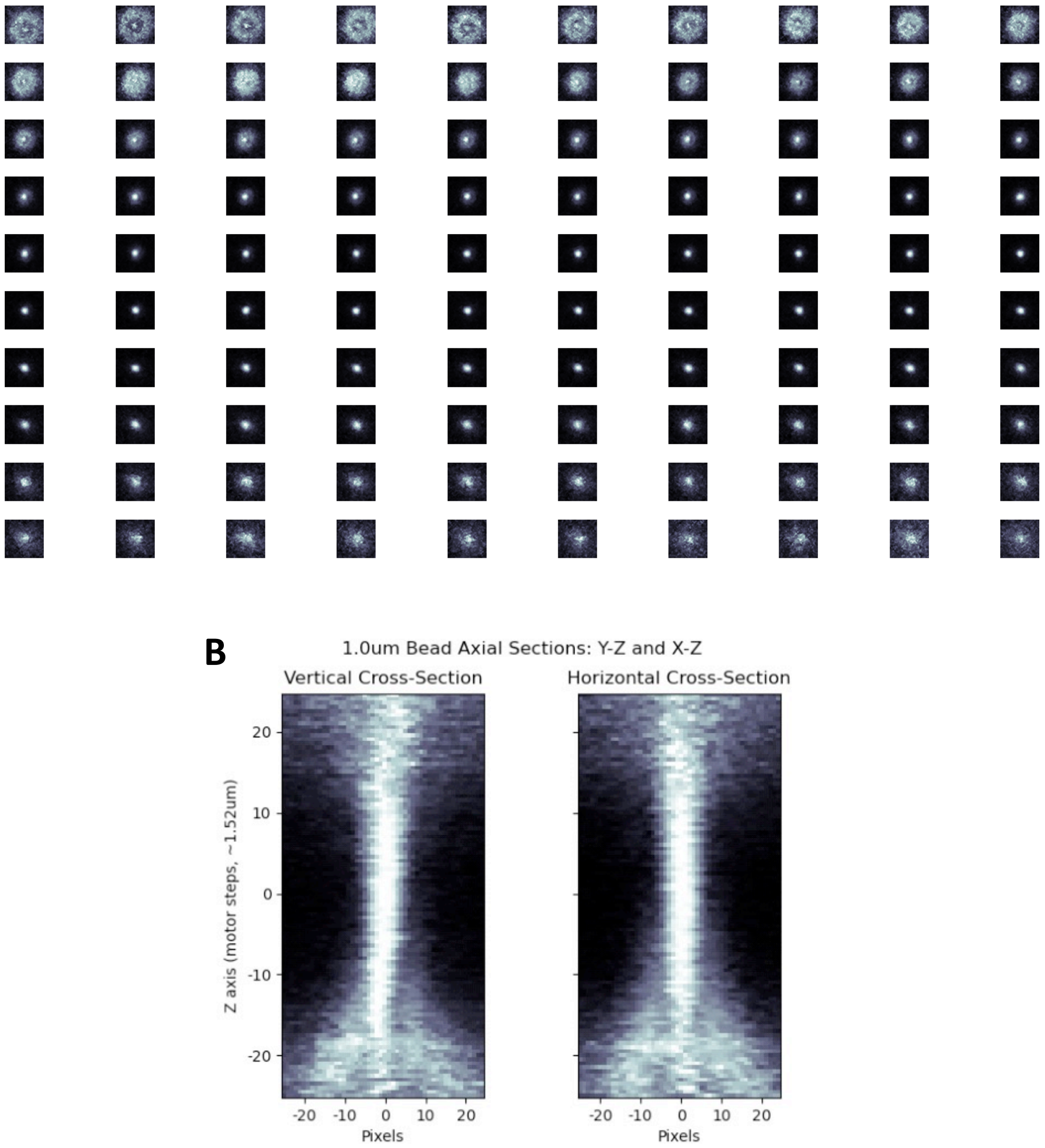


**Figure S6: (A)** Vertically sliced “Z-Stacks” of 1.0 μm polystyrene microspheres (calibration beads), captured by making single vertical steps for each slice. The object begins out of focus, comes into focus, and becomes out of focus as vertical space is traversed. **(B)** “Stacked” arrangement of the frames in (A), bisected to create a “cross-section” of the Z-Stack. The vertical distance within which the object remains in focus empirically describes the effective depth of focus of the optical system, approximately 20 steps at 1.52 μm per step corresponding to an effective 30 μm depth of focus.

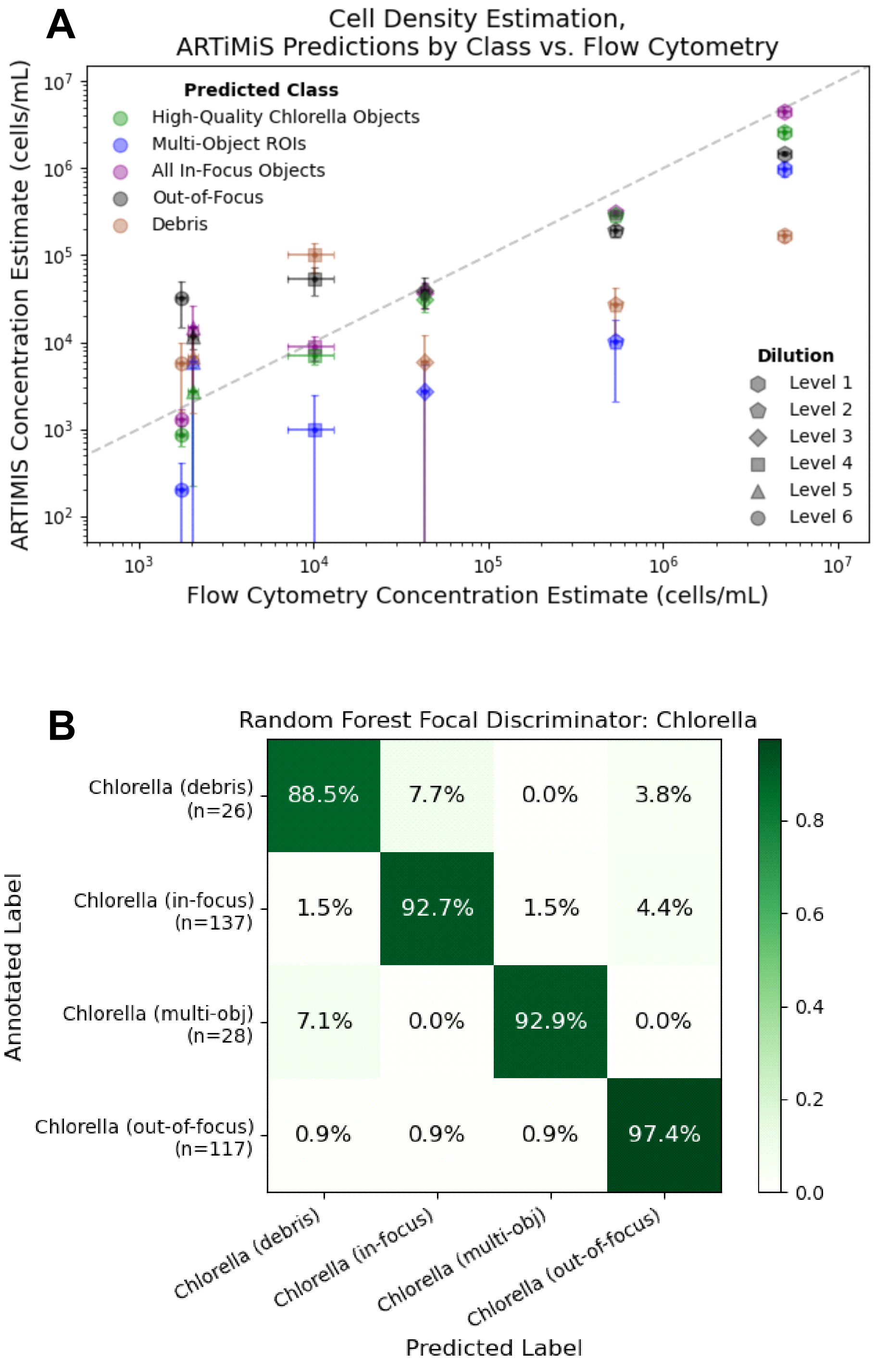


**Figure S7: (A)** Reproduced plot from Figure 4, including additional out-of-focus and debris classes which were omitted from Figure 4 for clarity. Concentration of debris was generally dilution-agnostic; out-of-focus particle concentration appeared to scale non-linearly with dilution, reaching a stationary threshold between $1x10^4$ to $4x10^4$ objects/mL. **(B)** Confusion matrix of the classification model used to estimate particle identity in sample points of (A).

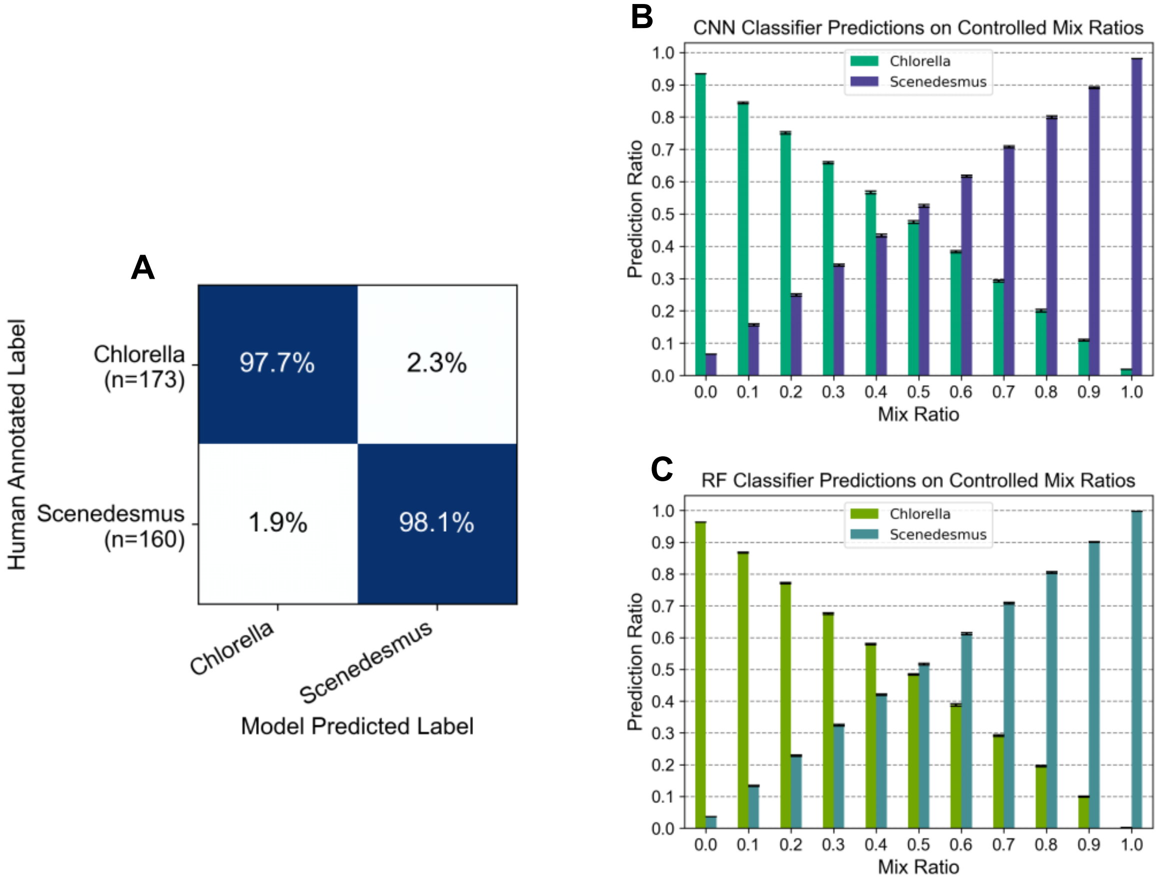


**Figure S8: (A)** Confusion matrix of convolutional neural network (CNN) model corresponding to random forest model described in Figure 5. **(B, C)** *In silico* mix ratio results when evaluating aggregate predictions at varying ratios of Class A to Class B; ratio of 0.0 corresponds to 100% *Chlorella* and 0% *Scenedesmus* in the test dataset; ratio of 1.0 corresponds to 0% *Chlorella* and 100% *Scenedesmus*. Unique particles in each dataset were randomly resampled 5 times, with bars depicting replicate means, error bars describing standard deviation. Panels contrast CNN classifier (B) to Random Forest classifier (C).

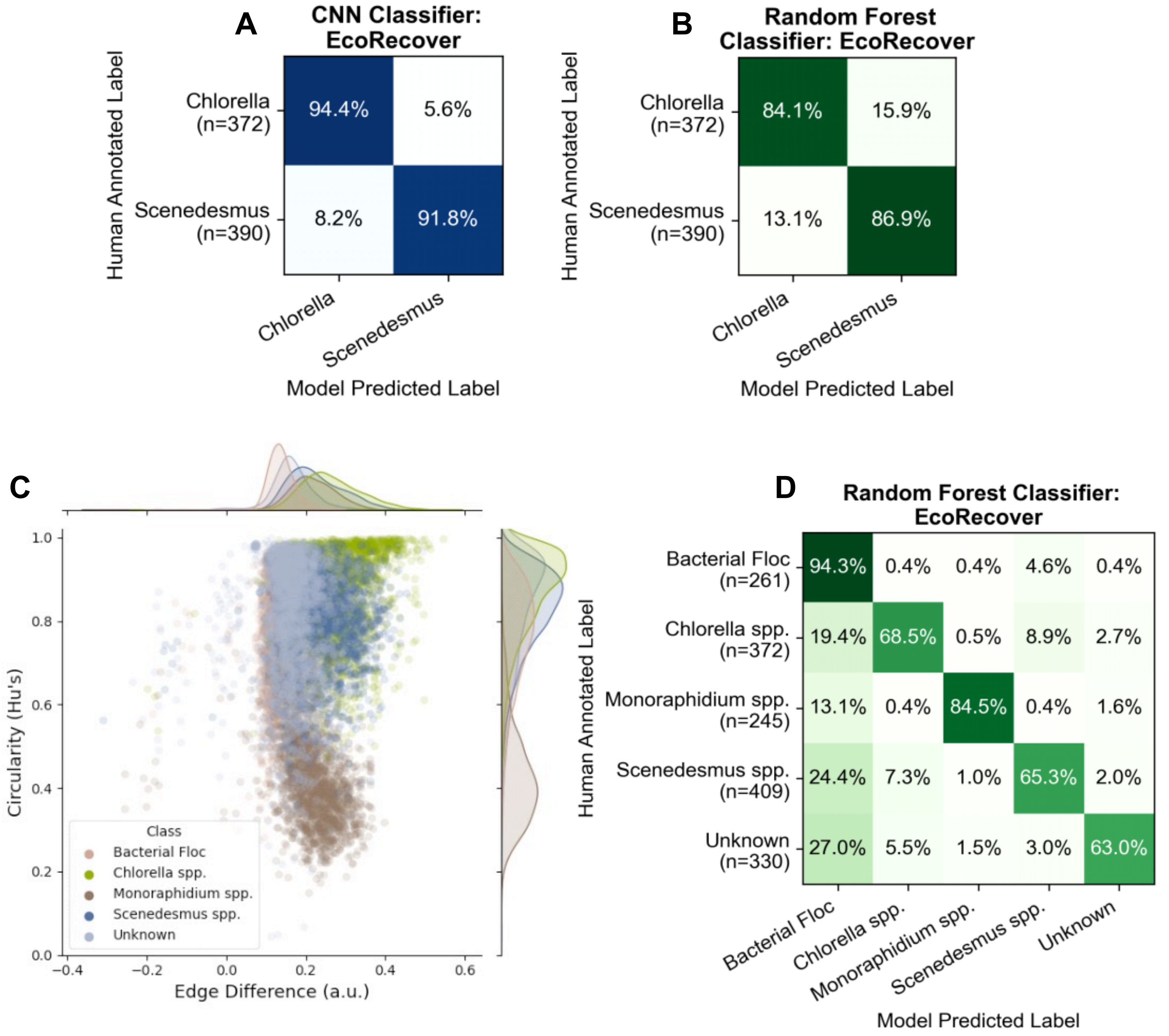


**Figure S9: (A)** CNN classifier trained to differentiate *Chlorella* from *Scenedesmus* using training and test data from the EcoRecover microalgal community dataset, and **(B)** random forest classifier trained on the same data. **(C)** Distribution of unique particles belonging to the dominant taxonomic groups found in the EcoRecover system, displaying two of the top important features for class differentiation derived from the random forest classifier in (D). **(D)** Confusion matrix of a grid-search optimized random forest classifier trained to distinguish between the classes described in (C) and Figure 6.
